# Characterizing vascular function in mouse models of Alzheimer’s disease, atherosclerosis, and mixed Alzheimer’s and atherosclerosis

**DOI:** 10.1101/2024.12.02.626393

**Authors:** Beth Eyre, Kira Shaw, Dave Drew, Alexandra Rayson, Osman Shabir, Llywelyn Lee, Sheila Francis, Jason Berwick, Clare Howarth

## Abstract

**Significance:** Alzheimer’s disease does not occur in isolation and there are many comorbidities associated with the disease – especially diseases of the vasculature. Atherosclerosis is a known risk factor for the subsequent development of Alzheimer’s disease, therefore understanding how both diseases interact will provide a greater understanding of co-morbid disease progression and aid the development of potential new treatments.

**Aim:** The current study characterizes hemodynamic responses and cognitive performance in APP/PS1 Alzheimer’s mice, atherosclerosis mice, and a mixed disease group (APP/PS1 & atherosclerosis) between the ages of 9 and 12 months.

**Approach:** Whisker-evoked hemodynamic responses and recognition memory were assessed in awake mice, immunohistochemistry to assess amyloid pathology, and histology to characterize atherosclerotic plaque load.

**Results:** We observed hemodynamic deficits in atherosclerosis mice (vs Alzheimer’s, mixed disease or wild-type mice), with reduced short-duration stimulus-evoked hemodynamic responses occurring when there was no concurrent locomotion during the stimulation period. Mixed Alzheimer’s and atherosclerosis models did not show differences in amyloid beta coverage in the cortex or hippocampus or atherosclerotic plaque burden in the aortic arch vs relevant Alzheimer’s or atherosclerosis controls. Consistent with the subtle vascular deficits and no pathology differences, we also observed no difference in performance on the novel object recognition task across groups.

**Conclusions:** These results emphasize the importance of experimental design for characterizing vascular function across disease groups, as locomotion and stimulus duration impacted the ability to detect differences between groups. Whilst atherosclerosis did reduce hemodynamic responses, these were recovered in the presence of co-occurring Alzheimer’s disease which may provide targets for future studies to explore the potentially contrasting vasodilatory mechanisms these diseases impact.

## 1 Introduction

Globally, there are over fifty-five million people living with dementia, with this set to increase to over one hundred and thirty-nine million by 2030^1^. Alzheimer’s disease (AD) is the most common cause of dementia, with hallmark features including the accumulation of proteins, such as amyloid beta plaques^2^ and hyperphosphorylated tau tangles^3^. Accumulating evidence suggests that neurovascular dysfunction may be important in the pathogenesis of AD; with vascular changes being observed as one of the first pathological changes in the disease^4^. According to the two-hit vascular hypothesis^5^ vascular risk factors, such as diabetes, hypertension, cardiovascular disease and/or cerebrovascular deficits result in damage to the blood-brain-barrier (BBB)^6,7^, resulting in hypoperfusion^8^. This damage to the BBB and reduction in blood flow to the brain alters the production and clearance of proteins such as amyloid beta in the brain^9,10^. An increased accumulation of proteins can be damaging to neurons, resulting in neuronal dysfunction, neurodegenerative changes and ultimately neuronal loss, leading to the cognitive impairment observed in AD.

Alzheimer’s is not a disease that occurs in isolation. As age is the greatest risk factor of the disease, many individuals possess comorbidities^11^. Individuals who possess a greater number of comorbidities have poorer outcomes^12^. AD and diseases of the vasculature share common risk factors, such as diabetes, hypertension and hypercholesterinaemia^13-16^. However, research assessing how vascular comorbidities impact neurovascular function are lacking. It is important to assess how comorbid disease may affect neurovascular function, especially within preclinical research as this could potentially explain why many animal studies assessing drug treatments for Alzheimer’s disease are unable to translate to humans^17-19^, for instance if they focus on singular mechanisms which are differentially modulated across disease types.

Atherosclerosis is the second biggest killer in the UK^20^, and is an inflammatory disease caused by an excess of lipids within the blood. Atherosclerosis can result in reduced perfusion, due to the occlusion of arteries, and many studies have observed associations between atherosclerosis and AD^21-23^. Atherosclerosis can be modeled preclinically in a number of ways^24^. For example, atherosclerosis genetic knockout mouse models such as the LDLR-/- ^25^ or APOE-/- ^26^ can be used. Neurovascular deficits have previously been reported in the LDLR-/- model of atherosclerosis, where Lu et al. (2019)^27^, reported weaker evoked-hemodynamic responses to a whisker stimulation, as well as hypoxic pockets in cortical tissue and microvascular changes in capillaries at 12 months of age. This group further observed that in the cortex of 12-month old atherosclerotic mice tissue oxygenation was reduced, red blood cell velocity and flux were slower, and capillary diameters smaller^28^. However, breeding genetically modified mice can be time consuming and expensive, especially when wanting to model atherosclerosis concurrently with other diseases. Recently, other ways of inducing atherosclerosis have been employed, via the use of viral vectors^29-31^. Injecting a gain-of-function mutation of PCSK9 via a viral vector (in addition to a western diet) can increase cholesterol levels^29,30^ and over time result in atherosclerotic lesions. Using this induced model, recent research has investigated whether atherosclerosis alone and in the presence of amyloid overexpression can impact neurovascular function in the J20-AD mouse model in a lightly anesthetized preparation^31^. Shabir et al., observed that the presence of atherosclerosis resulted in a reduction in the size of the peak hemodynamic response to a 2s whisker stimulation. However, they observed no effect of AD alone or mixed Alzheimer’s and atherosclerosis on evoked-hemodynamic responses, which they speculated could be due to the presence of amyloid plaques invoking compensatory angiogenesis in cerebral microvessels to enhance perfusion in response to increased Aβ and neuroinflammation.

In this study we combined comorbid Alzheimer’s and atherosclerosis using a viral injection of a gain-of-function mutation of PCSK9 (with a western diet) in a more ‘severe’ Alzheimer’s disease model, the APP/PS1 mouse strain^32,33^. Whilst a previous study has used this same method to induce hypercholesteremia in the APP/PS1 model of AD^30^, they only investigated the impact of mixed disease on amyloid plaque number in the hippocampus. Therefore, in this novel investigation we aimed to assess how AD alone, atherosclerosis alone, and mixed AD and atherosclerosis impact recognition memory, amyloid pathology and sensory-induced vascular function in the awake mouse^34^.

## 2 Methods

### 2.1 Animals

Male mice aged between 9-12 months from the following groups were used: APP/PS1 (B6.C3-Tg(APPswe,PSEN1dE9)85Dbo/Mmjax #34829) Alzheimer’s model^35^, referred to as AD throughout the manuscript; wild-type littermates referred to as WT, an atherosclerosis model (WT-littermates injected with rAAV8-mPCSK9-D377Y (6 × 10^12^ virus molecules/ml) (Vector Core, Chapel Hill, NC) at 11 weeks of age (i/v or i/p + a western diet at 12 weeks (21% fat, 0.15% cholesterol, 0.03% cholate, 0.296% sodium; #829100, Special Diet Services UK)) referred to as ATH; and a mixed disease group (APP/PS1 mice injected with rAAV8-mPCSK9-D377Y (6 × 10^12^ virus molecules/ml) (Vector Core, Chapel Hill, NC) at 11 weeks of age (i/v or i/p + a western diet at 12 weeks)) referred to as MIX. Western diet began at 12 weeks and continued until the end of the study. Prior to surgery, mice were housed in litters where possible, however some were singly house for welfare reasons or if there were no available littermates. A 12hr dark/light cycle (lights on 06:00-18:00) was implemented. Food and water were accessible ad-libitum with the western diet limited to 5g per day per mouse. Experiments were completed during the light cycle, between the hours of 8am and 4pm.

Only male mice were included in the current manuscript. For the two groups with induced atherosclerosis, we only collected data from male mice due to differential expression of mPCSK9 in the livers of males and females^36^. For the remaining two groups where data were collected from male and female mice (WT, APP/PS1), we observed sex differences in stimulus-induced hemodynamic responses and performance on the novel object recognition task (see Supplementary Figures 1-3)- which instead will be explored further in a separate more in-depth investigation.

Procedures used in the study were approved by the UK Home Office and in agreement with the guidelines and scientific regulations of the Animals (Scientific Procedures) Act 1986. Further approval was also granted by the University of Sheffield licensing committee and ethical review board. The following study is reported in accordance with the ARRIVE guidelines. Once mice with the appropriate date of birth were identified, selection was randomized. The experimenter was blinded from the disease group where possible during both the collection and analysis. A priori sample sizes were not conducted.

### 2.2 Novel Object Recognition Task

At 9 months of age non-spatial recognition memory was assessed using the novel object recognition (NOR) test (Figure 1a; Figure 5). A 2-day protocol was used based upon the protocol developed by Lueptow (2017)^37^. On day 1 the habituation phase took place. Mice were placed into a square open field arena (40x40x40cm) for 10 minutes and behavior was recorded using a camera placed above the arena. Day 2 consisted of the training and testing phases. In the training phase mice were placed into the open field arena with two identical objects (either two glass beakers or two Duplo tower blocks – these were counterbalanced across mice). Objects were placed diagonally from each other; positioning of objects was also counterbalanced across mice. After 10 minutes mice were removed from the arena. After a 1-hour retention period mice were placed back into the arena. However, one of the familiar objects had been replaced with the novel object. Mice were handled for approximately 7 days prior to cognitive testing to reduce experimental stress. The arena and objects were cleaned with 70% ethanol between mice. At least 30 minutes before cognitive testing mice were moved to the experimental room to habituate them to the new room and to reduce stress. Time spent with each of the objects was measured using EthoVision software (EthoVision XT 15, Noldus). Mice were deemed to be exploring an object if their nose was <2cm away from the object. However, climbing on top of objects was not considered exploration and was not included as exploration of objects. Distance traveled and velocity during training and testing was also recorded. Exploration of individual objects allowed for the calculation of the preference index. Preference index was calculated by dividing the time spent with the novel object with the total exploration of both objects multiplied by one hundred (during the testing phase). This produced a percentage score of preference for the novel object, with a score of 50% suggesting no preference for the novel object, a score >50% indicating a preference for the novel object and a score <50% indicating a preference for the familiar object. To be included in the analysis mice had to explore both familiar objects (in the training phase) for a minimum of 20s each.

**Fig. 1.**
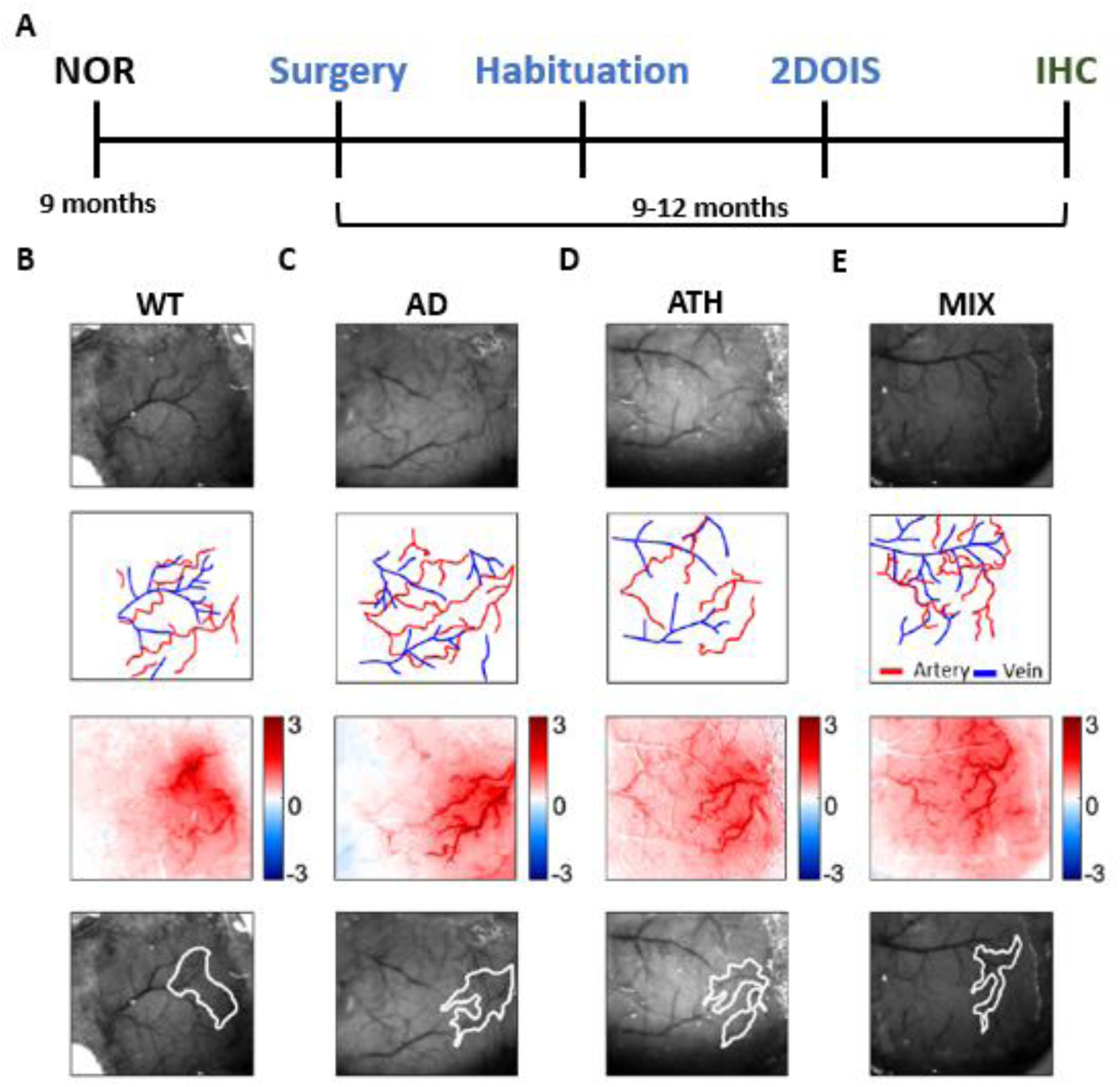
Experimental timeline and representative 2D-OIS spatial images. **A.** Experimental timeline for all mice included in these experiments. Subjects first underwent cognitive testing on the novel object recognition (NOR) task at 9 months, before a surgery to insert a headplate and thinned window over the whisker barrel cortex was conducted. After a recovery period (minimum 1 week) to the imaging apparatus (5 days), regional hemodynamic responses were recorded in awake mice using 2-dimensional optical imaging spectroscopy (2DOIS). Mice were sacrificed using transcardial perfusion, and immunohistochemical (IHC) staining was conducted to assess amyloid coverage in AD and MIX groups. Histology was also conducted on the ATH and MIX groups to assess atherosclerotic plaque burden in the aortic arch. Representative animal from each disease group **B.** WT **C.** AD **D.** ATH and **E.** MIX revealing the thinned window region (row one), vessel maps revealing arteries and veins (row two), a spatial map of HbT changes to a 2s whisker stimulation (row three) with red colors indicating an increase in HbT in response to a 2s whisker stimulation, and the region of interest overlying the active whisker barrel region (row four, white ROI).

### 2.3 Surgery

Surgery to implant a head plate and thinned cranial window was conducted between 9-12 months of age (Figure 1a). Briefly, anesthesia was induced using ketamine (50mg/kg) and medetomidine (0.65mg/kg) (subcutaneously injected, s/c). The surgical plane of anesthesia was maintained with the addition of isoflurane (0.5-0.8% in 100% oxygen). Carprofen (10mg/kg, s/c) was administered prior to removal of hair from the head. Animals were placed in a stereotaxic frame (Kopf Instruments) and ophthalmic gel was administered (Viscotears, Novartis). Body temperature was constantly monitored and sustained using a rectal thermometer and a homeothermic blanket (Harvard Apparatus). Iodine and bupivacaine (50-100mcL at 0.025%) were applied prior to exposing the skull. Suture lines were covered with cyanoacrylate glue and a dental scraper scored the contralateral side of the skull. The bone overlying the right somatosensory cortex was thinned to translucency (∼4mm^2^) using a dental drill. Saline was administered throughout to cool the area and to assist with the visualization of the pial vasculature. Cyanoacrylate glue was applied to the thinned region. The metal head plate was attached using dental cement (Superbond C & B; Sun Medical). Atipamezole (2mg/kg in 0.3ml warm sterile saline s/c) was administered at the end of the procedure to reverse the effects of medetomidine. Following surgery, mice were placed in an incubator (29 degrees C) and monitored. After surgery, mice were given at least one week to recover prior to habituation and awake imaging. Mice were closely observed for weight loss and signs of pain for 3-days after surgery and administered carprofen jelly (10mg/kg) for at least 1-day.

### 2.4 2-Dimensional Optical Imaging Spectroscopy (2D-OIS) in Awake Mice

After a minimum 1-week post-surgical recovery period mice underwent hemodynamic imaging using 2D-OIS at 9-12 months (Figure 1a). The awake imaging set up was previously described in Eyre et al. (2022)^38^, however a modified habituation and imaging procedure was used in the current study. Briefly, one week after surgery mice were habituated to the awake imaging spherical treadmill set-up. Over 5 days, mice were familiarized to the experimental room, experimenter and awake imaging apparatus. On day one, mice were placed on the spherical treadmill for 10 minutes, the room light was kept on, and mice were not head-fixed. Day 2, mice were head-fixed and placed on the spherical treadmill, lights were turned off and 2s whisker stimulations were conducted using a mechanical plastic T-bar. Mice were on the ball for approximately 20 minutes. Day 3 was the same as day 2, with a supplementary ‘spontaneous’ experiment conducted, where hemodynamic measurements were collected but no whisker stimulation applied. Mice were on the ball for approximately 30 minutes. Day 4 followed the same procedure as the preceding two days; however, an additional 16s whisker stimulation experiment was conducted. Mice were on the ball for approximately 45 minutes to 1 hour. Day 5 followed the exact same procedure as day 4. Mice received a reward of sunflower seeds following each experimental imaging day. Hemodynamic data was collected on all days where mice were head-fixed and used in the subsequent analysis (Figure 1b-e). Mice were briefly anesthetized using isoflurane (3-4%) prior to being placed on the awake imaging apparatus.

Locomotion behaviors were gathered using a spherical treadmill with an optical motion sensor. In-house MATLAB scripts (MathWorks, 2024a) were used to analyze the locomotion data. The optical motion sensor logged treadmill movement during all experiments. Locomotion data files were composed of: the locomotion data (a vector with zeros when the mouse was stationary and integers (arbitrary units) when the mouse was moving); a corresponding time vector (to capture frames per second); and the trigger points which specified the explicit timing of whisker stimulations across trials for aligning across imaging modalities.

Changes in hemoglobin concentration to a 2s or 16s whisker stimulation were investigated in the surface vasculature using widefield imaging. Two-dimensional optical imaging (2D-OIS) uses four wavelengths of light to measure changes in oxygenated (HbO), deoxygenated (HbR) and total levels of hemoglobin (HbT). Cortical hemodynamics were investigated using 4 differential wavelengths of light (494 ± 20 nm, 560 ± 5 nm, 575 ± 14 nm and 595 ± 5 nm). These wavelengths illuminated the thinned window region of the cortex (Figure 1b-e, row one), using a Lambda DG-4 high-speed galvanometer (Sutter Instrument Company, USA). A Dalsa 1M60 CCD camera was implemented to capture remitted light at 184 × 184 pixels, at a 32 Hz frame rate, providing a resolution of ∼75µm. 2-D spatial maps of micromolar changes in HbO, HbR and HbT were collected which reveal changes in hemoglobin concentrations in the surface vasculature (Figure 1b-e, row three). This is accomplished by implementing a path length scale algorithm (PLSA) to complete a spectral analysis. The PLSA uses the modified Beer Lambert Law, with a path-length correction factor, in addition to predicted absorption values of HbT, HbO and HbR^31,39,40^. Relative concentration estimates of the above were obtained from baseline values, where the hemoglobin concentration within the tissue was estimated as 100 µM, and tissue saturation of oxygen within the whisker region estimated at 80% and artery region at 90%.

Regions of interest (ROIs) were generated from the previously described 2D spatial maps, using in-house MATLAB scripts^31,39,40^. A whisker region was made using code that found the region of the cortex with the greatest change in HbT to a 2s whisker stimulation. Pixels were considered ‘active’ if they had a value that was >1.5 standard deviation (STD) across the entire spatial map of the surface vasculature. The whisker ROI was therefore the region of the cortex in which there was the largest increase in HbT in response to a 2s whisker stimulation (Figure 1b-e, row four). ROIs for an artery, vein and parenchyma were also manually selected within the whisker region. ROIs were generated for each imaging session. Care was taken to select the same artery, vein and parenchyma across imaging sessions for the same mice. Time series analyses in the current study were conducted on the active artery from within the whisker barrels (although are visualized for the overall whisker region, or the vein or parenchyma within the whisker barrels in Supplementary Figures 4-5).

### 2.5 Hemodynamic Data Analysis

For 2 and 16s-whisker stimulation experiments, hemodynamic responses (HbO, Hbr, HbT) were cut into trials around the stimulation period. For 2s experiments these trials started 5s before and 20s after the onset of the stimulation, and for 16s experiments 10s before and 60s after the onset of the stimulation. All 2s stimulation experiments consisted of 30 trials, and 16s experiments of 15 trials. Each trial was extracted individually alongside its corresponding locomotion trace. The locomotion traces were then classified as belonging to one of four categories: 1- no running during the stimulation, 2- running before but not during the stimulation, 3- running starts at onset of stimulation, or 4- running before and during the stimulation. For this locomotion classification across trials the running traces for the 4s before and 4s after stimulus onset/offset were assessed. For each individual trial the area under the curve (AUC) of the hemodynamic and locomotion responses were assessed during and just beyond the stimulation period (5-10 seconds for 2s stimulation or 10-30 seconds for 16s stimulation) using the ‘trapz’ function in MATLAB, and the maximum peak by finding the largest value reached during this period. These time series metrics and locomotion group categorizations were then extracted in a table alongside the animal ID, disease group (WT, AD, ATH or MIX), imaging session ID (day 1-4) and trial ID (1-30 for 2s, 1- 15 for 16s) labels. Tables were then exported into R, where statistical analysis was conducted on individual trials using a linear mixed model. Only stimulation trials where there was no running (locomotion group 1) or where running started at the onset of the stimulation (locomotion group 3) were included in the main analysis due to the profound impact of locomotion on hemodynamic responses. In supplementary figures expanding the investigation of the impact of locomotion on hemodynamic responses, all experimental trials were included and ranked in ascending order from least to most locomotion for the 5 seconds after stimulation onset (5-10 seconds) using the locomotion AUC metric previously described (Supplementary Figure 7).

### 2.6 Histology and Immunohistochemistry

At the end of the final experiment, mice were euthanized with pentobarbital (100 mg/kg, Euthatal, Merial Animal Health Ltd). Cardiac perfusions were completed using saline (0.9%). A subset of mice were also perfused with formalin (10%). Brains were dissected and the whole brain (or half) was either placed in formalin (10%) or snap frozen in isopentane and stored in a -80C freezer. Formalin fixed brains were paraffin embedded (FFPE) and were cut into coronal slices (5-7µm) using a vibratome. Coronal sections were mounted onto slides for subsequent immunohistochemistry. An avidin-biotin complex (ABC) method was used to stain and quantify amyloid coverage. Briefly, slides were dewaxed with xylene and rehydrated using ethanol (100%, 100%, 95%, 70%) for 5 minutes each. Peroxidase activity was inhibited by placing in a bath of 3% H_2_O_2_/12ml methanol for 20 minutes. Following this, slides were placed in 70% formic acid for 10 minutes. Antigen retrieval was completed using a microwave oven in a buffer of trisodium citrate (PH6) for 10 minutes. After antigen retrieval, slides were placed in a bath of dH_2_O for 10 minutes. Slides were washed in PBS (for 5 minutes x2). Slides were then incubated with 1.5% normal serum (goat) for 30 minutes at room temperature and then incubated with the primary antibody (Amyloid Beta Rabbit Monoclonal – 1:500, Abcam, ab201060) overnight at 4 degrees C. Slides were again washed in PBS (for 5 minutes x2) and the secondary antibody was applied for 30 minutes at room temperature. Slides were then washed with PBS (for 5 minutes x2). A horseradish peroxidase avidin-biotin complex (Vectastain Elite Kit, Vector Laboratories, UK) was then applied and left at room temperature for 30 minutes. A final wash in PBS (for 5 minutes x2) was completed and then 3,3-diaminobenzidine tetrahydrochloride (DAB) (Vector Laboratories, UK) was applied to help visualize the antibody (left for 3 minutes). Slides were rinsed with dH_2_O for 3-5 minutes and counterstained with haematoxylin and differentiated with Scotts Tap Water. Finally, slides were dehydrated using baths of ethanol (70%, 90%, 100%, 100%) for 30 seconds each and placed in xylene for 2-3 minutes. Following dehydration, slices were mounted with coverslips using DPX. Stained slides were scanned using a slide scanner (3Dhistech Panoramic 250) at a 40x magnification.

A classifier model in Qupath was used to assess amyloid burden within brain tissue^41^. To create the classifier model, a high resolution (0.88*μ*m/pixel) random trees pixel classifier was trained on a subset of stained brain sections. For the cortex ROI, scanned images were viewed using the haematoxylin channel and a rectangle ROI was drawn on an area of the cortex above the hippocampus. For the hippocampus ROI the DAB channel remained on and an ROI was drawn around the entire hippocampus. The classifier was applied and then a manual inspection was conducted. Upon manual inspection, if areas of tissue outside the ROI were included these were reclassified as ‘ignore’. Additionally, if cerebral amyloid angiopathy (CAA) was classified as plaque, this was manually reclassified as ‘CAA’ and any CAA not detected by the model was manually added. If poorly perfused vessels were classified as amyloid, these were reclassified as ‘tissue’. Amyloid burden included both amyloid plaques and CAA, and this was calculated by the following formula:

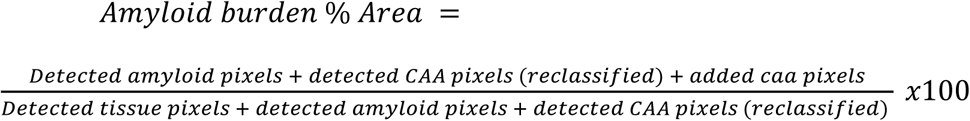

### 2.7 Identification and quantification of atherosclerosis

After cardiac perfusion, the heart and aorta were also dissected for atherosclerotic and mixed disease mice. The heart and aorta were blunt dissected and placed in formalin for a minimum of 24 hours before being placed in PBS. They remained in PBS until they were stained and embedded into wax filled petri dishes. Visible fat was removed from the aorta under a dissecting microscope and the aorta was cut from the heart at the aortic root. Dissecting scissors were used to cut the aorta down the middle to open the aortic arch. This allowed the aorta to be laid flat and pinned to wax in a petri dish post-staining. Oil red O (Sigma, O0625-100G) was used to stain for natural triglycerides and lipids within the aortic arch. A solution of Oil red O (60% solution) was made using distilled water and isopropanol. Once the aorta had been cut it was stained. Briefly, the aorta was placed into distilled water (10 seconds) then isopropanol (2 minutes) and then into the Oil Red O solution (6 minutes). It was then placed in isopropanol (2 minutes) and finally rinsed in distilled water. Following staining, petri dishes were filled with wax. Once the wax had partially hardened 1mm insect pins (Fine Science Tools, FST) were used to pin the edges of the aorta into the wax. A 12MP camera, placed 10 cm above the samples was used to take images of stained aortas. ImageJ was used to quantify the presence of atherosclerotic plaque burden in the aortic arch. The polygon tool was used to draw around the aortic arch (the ROI) and the area of this was measured. Images were converted to 8-bit grayscale images. A threshold was applied to identify positively stained Oil Red O areas. A threshold of 150 was applied to all samples. Percentage area of aortic arch plaque burden was calculated using the following equation:

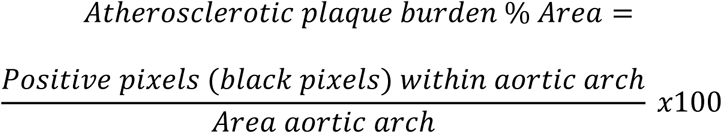

Two aortas were excluded from the analysis due to significant damage to the aortic arch during the dissection.

### 2.8 Statistical Tests

Statistical tests were conducted in RStudio and GraphPad Prism (Version 10), and figures created in GraphPad Prism and MATLAB (R2024a). The threshold for statistical significance was set at p ≤ 0.05. All data are presented as mean ± SEM or individual data points unless otherwise stated. The number of animals and experimental trials are reported across all comparisons. Detailed statistical reports are included in the supplementary information (Tables SR1-SR6).

Multi-group comparisons (across disease group (WT, APP, ATH, MIX) only, or disease group and trial number (1-30, 1-15) or sex (M, F) interactions) were conducted using linear mixed models, with animal ID inputted as the random factor to account for variations between groups being driven by a single outlier animal (lmer package RStudio). For behavioral data analysis (Figure 5) where there was only one independent variable (disease group) and each animal only contributed one datapoint, a one-way ANOVA was conducted across all metrics (distance traveled, velocity, preference index) as data was normally distributed within each group (assessed using a Shapiro-Wilks test). Where post-hoc comparisons were conducted to explore significant effects, the Tukey method was used with correction for multiple comparisons. For correlational analysis exploring the relationship between the trial number or size of locomotion and hemodynamic responses (Figure 3; Supplementary Figure 6) a Pearson’s R correlation was conducted to assess whether the variables were significantly linearly related, and to compare the linear relationship between the disease groups a linear regression was conducted to compare the slope and intercepts.

## 3 Results

### 3.1 Atherosclerosis mice show a reduced hemodynamic response to a short 2s stimulation, but not a longer 16s stimulation

We compared short (2s) and long (16s) duration stimulation-induced hemodynamic responses from the artery within the whisker barrel region of somatosensory cortex across disease groups in rest trials where locomotion did not occur during the stimulation period (Figure 2; Statistics Report 1). In response to a 2s-whisker stimulation (Figure 2A) there was a significant effect of disease on the maximum peak of the HbT trace (p=0.008; Figure 2B)) and the area under the curve of the HbR trace (p=0.03; Figure 2C) as the atherosclerosis mice showed smaller hemodynamic responses than those in WT mice (WT vs ATH: HbT peak p =0.02; HbR AUC p=0.01). Of particular interest, the size of the HbT peak in atherosclerosis mice was significantly improved in the mixed model (ATH vs MIX p=0.03), replicating the findings of Shabir et al., who used the J20 model. These differences were not driven by differences in locomotion, as the confounds of locomotion were removed from this comparison by selecting only trials with no concurrent locomotion during the stimulation period (effect of disease on size of locomotion events: p=0.31; Figure 2D). In contrast, there was no significant effect of disease group on the hemodynamic response induced by a 16s whisker stimulation (Figure 2E), as the size of the HbT maximum peak (Figure 2F) and HbT area under the curve (linear mixed model: F=0.1567, p=0.92; data not shown) was comparable between groups. Although no overall significant difference was observed in the size of the 16s stimulus-induced HbR response, there was a trend level effect of disease (p=0.06; Figure 2G), as the mixed model showed a smaller HbR washout versus the wild-type animals. Again, locomotion did not differentially influence hemodynamic responses in the 16s stimulation dataset as there was no difference between groups in the amount of locomotion during the stimulation period due to only rest trials being included (Figure 2H). Our results were replicated when we assessed 2s and 16s stimulation-evoked hemodynamic responses in our other regions of interest (the entire whisker barrel region, vein, and parenchyma) (Supplementary Figures 4-5).

**Fig. 2.**
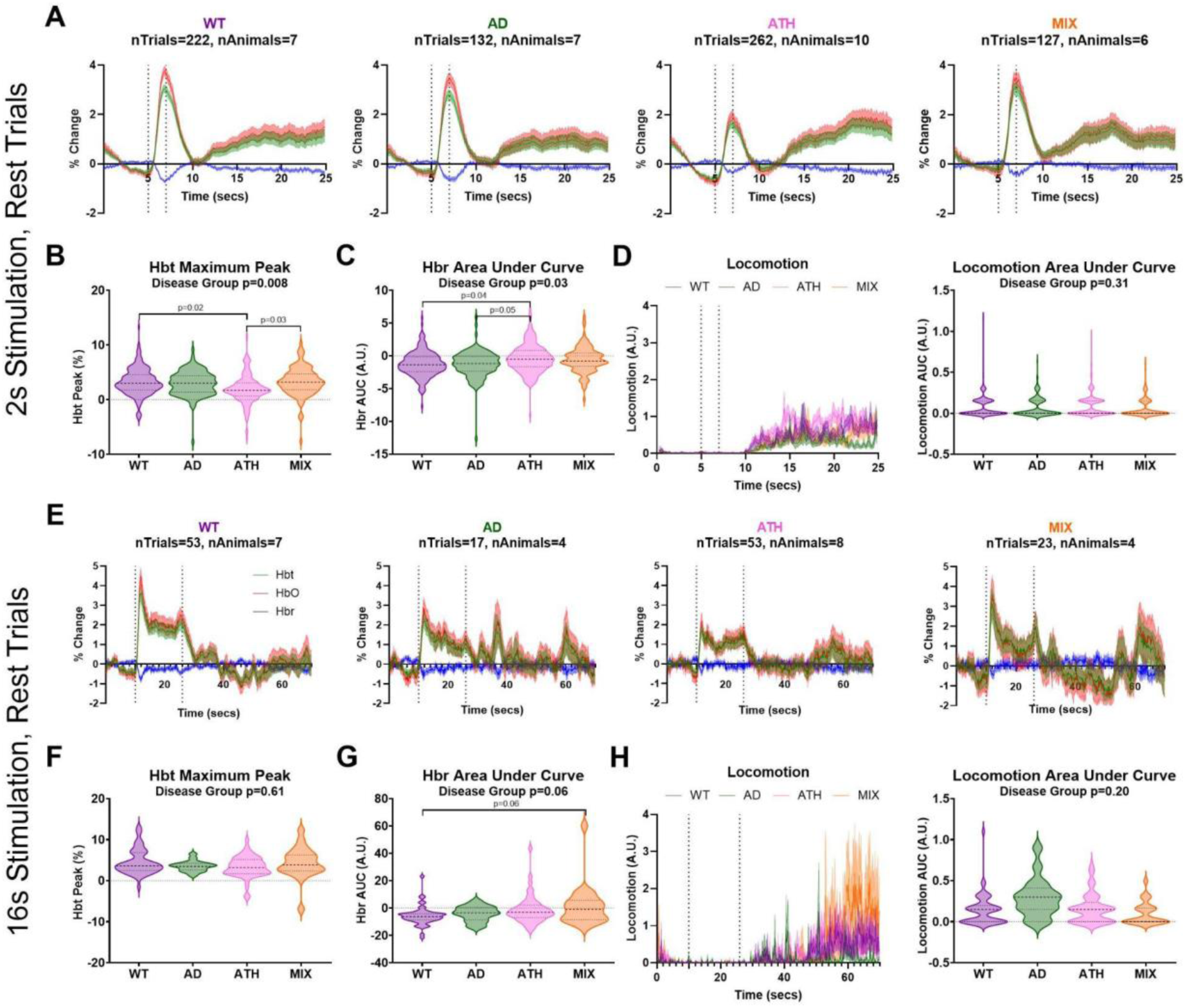
Comparing stimulation-induced responses without the confounds of locomotion. **A.** Hemodynamic time series from the artery ROI within the whisker barrel region showing total (HbT, green), oxygenated (HbO, red), and deoxygenated (HbR, blue) hemoglobin in response to a 2s mechanical whisker stimulation in trials with no concurrent locomotion occurring between the 4 seconds either side of the stimulus period (dotted lines) for wild-type (WT, purple), APP/PS1 (AD, green), atherosclerosis (ATH, pink), and mixed APP/PS1 x atherosclerosis (MIX, orange) mice. **B.** The maximum peak of the HbT response during the stimulation period of these ‘rest trials’ is compared across disease groups using a linear mixed model, with a significant overall effect of disease (p=0.008) driven by the atherosclerosis group (pink) showing smaller 2s-stimulus induced responses than the wildtype (purple, p=0.02) or mixed (orange, p=0.03) mice. **C.** There was a significant difference between disease groups on the area under the curve of the HbR response during the stimulation period of these ‘rest trials’ (p=0.03), driven by the atherosclerosis group showing smaller HbR responses than the wild-types (p=0.01) and AD mice (p=0.05). **D.** There were no differences in locomotion during these ‘rest trials’, assessed using the area under the curve of the locomotion response during the stimulation period (right). **E.** Hemodynamic responses were also compared in response to a 16s mechanical whisker stimulation in trials with no concurrent locomotion occurring between the 4 seconds either side of the stimulation period (dotted lines) for the artery ROI. There was no significant difference in the size of the **F.** HbT (maximum peak, p=0.61) or **G.** HbR (AUC, p=0.06) responses between WT, AD, ATH or MIX mice, although the HbR washout was smaller at trend level due to the mixed APP/PS1 x atherosclerosis mice showing smaller responses than wild-type. **H.** There were also no differences in locomotion during rest trials for 16s stimulation events. P-values are taken from linear mixed-effects models with disease group inputted as fixed-effect factor, and animal ID as the random effect (lmer package RStudio), and pairwise comparisons (with correction for multiple comparisons) conducted using the Tukey method (emmeans package RStudio). Shaded error bars represent mean +/- SEM. Horizontal lines on violin plots show median and interquartile range. The number of trials and animals included in each group are indicated on the time series graphs.

### 3.2 The trial number does not influence stimulus-evoked hemodynamic responses

Given we observed differences between disease groups in the hemodynamic responses to a 2s whisker stimulation but not to a 16s whisker stimulation, we wondered whether the higher number of trials in the 2s experimental paradigm (30 trials vs 15 trials for 16s paradigm) could be driving our differences, as for instance the atherosclerosis group may be showing reduced hemodynamic activity only in later trials where the disease could be impacting the ability to sustain functional hyperemia (as seen in early aging by Balbi et al., 2015^42^). As such, we investigated whether the trial number (with higher numbers indicating trials which occurred later in the experiment) impacted the size of hemodynamic responses to a 2s stimulation (Figure 3; SR2-3). We showed no differences in the relationship between the size of the HbT response (assessed using maximum peak) and the trial number across disease groups (p=0.15; linear regression analysis), although when HbT peak and trial number were correlated within each individual disease group the wild-type animals showed significantly increased HbT responses to later trials (p=0.02) where the other groups did not (individual results in figure legend). We also saw a significant difference in the intercept of the correlation line between disease groups (p<0.0001), as the atherosclerosis group intercepted the y axis at a lower value, indicative of the smaller HbT response observed for this group (Figure 3A). To further explore the relationship between the size of the hemodynamic response and the trial number we also visualized the average HbT traces for early (1-5) vs late (25-30) trials across disease groups, and saw no effect of disease or trial number on the size of the HbT maximum peak, although there was a trend level interaction between disease group and trial number (p=0.07), driven by wild-type mice showing larger HbT responses in later trials (Figure 2B-C). The reason the overall effect of disease was lost in this comparison was likely due to the lower power to detect differences in this comparison due to the lower number of trials included (e.g. WT animals had 22 trials for the early group and 55 for the late group, versus 222 trials in the overall comparison for Figure 2, Figure 3A and Supplementary Figure 4). When we conducted the same analysis on the HbR response using area under the curve (Figure 3D-E) we did see a relationship between the size of the HbR response and the trial number for the mixed (atherosclerosis x Alzheimer’s) group (p=0.009) but not the other groups (significance values for individual groups reported in figure legend), but no overall relationship between trial number and size of the HbR response across groups (p=0.34). Again there was an overall significant difference in the intercept of the HbR AUC between groups, as the atherosclerosis group showed smaller responses (p<0.0001) (Figure 3D). When the average HbR traces were visualized for early (1-5) and late (25-30) trials (Figure 3E-F) there was no longer an overall impact of disease on the size of these responses, but there was a trend-level impact of trial number with the later trials showing larger decreases in HbR during the stimulation (p=0.07). The loss of the overall significant effect of disease in this comparison was again likely driven by the lower number of trials available reducing the statistical power for detecting differences between groups.

**Fig. 3.**
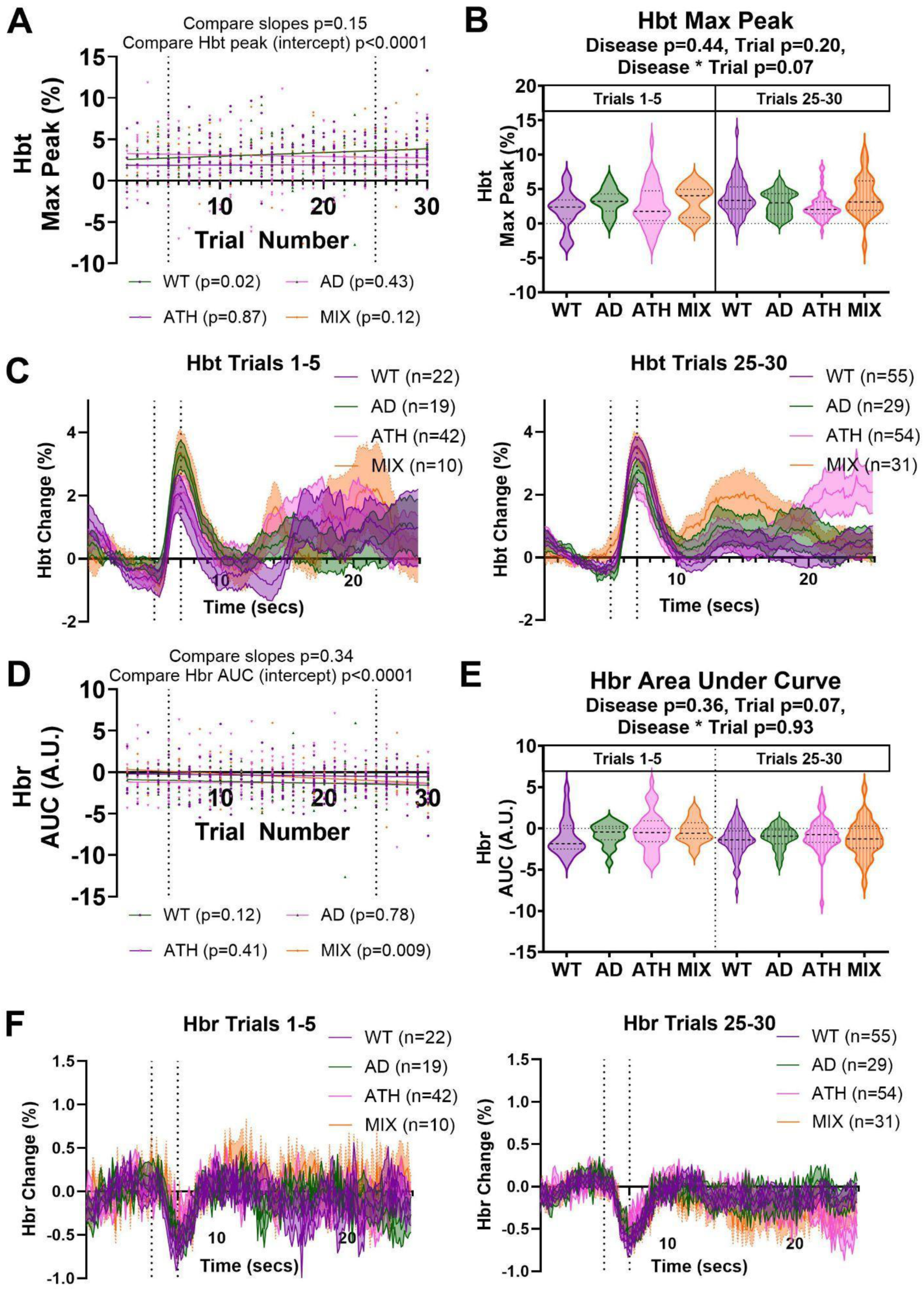
Differences across disease models in the size of 2s-stimulation responses during rest are not linked to the high number of trials. **A.** There was no correlation between the trial number (1-30) and size of the HbT response (max peak) for any of the disease groups (WT (p=0.22), purple; AD (p=0.38), green; ATH (p=0.59), pink; MIX (p=0.14), orange), and no significant impact of disease on the correlation between these two measures (compare slopes, p=0.36). However, there was an overall significant difference in the intercept (size of HbT peak) between disease groups (p<0.0001), with atherosclerosis mice consistently showing lower HbT peak values across all trials. **B-C.** When HbT responses were separated as belonging to early (trials 1-5) or late (trials 25-30) trials and compared across disease and trial groups there were no significant differences found. **D.** There was also no correlation between the trial number (1-30) and size of the HbR response (AUC) for most of the disease groups (WT (p=0.26), purple; AD (p=0.76), green; ATH (p=0.12), pink), although the MIX mice did show larger HbR responses in later trials (p=0.01, orange). Overall, there was no significant impact of disease on the correlation between these two measures (compare slopes, p=0.26), however, there was an overall significant difference in the intercept (size of HbR AUC) between disease groups (p<0.0001), with WT mice consistently showing larger HbR AUC values across all trials. **E-F.** When HbR responses were separated as belonging to early (trials 1-5) or late (trials 25-30) trials and compared across disease and trial groups, a significant effect of trial was observed as HbR responses were larger in later trials. On scatterplots individual values (dots) represent single trials, and p-values the result of a simple linear regression to test the correlation between variables within each disease group (legend) or the difference between groups on the slope and intercept (title). Shaded error bars represent mean +/- SEM. Horizontal lines on violin plots show median and interquartile range. P-values on violin plots are taken from linear mixed-effects models with disease and trial group inputted as fixed-effect factors, and animal ID as the random effect (lmer package RStudio), and pairwise comparisons (with correction for multiple comparisons) conducted using the Tukey method (emmeans package RStudio).

### 3.3 There are no longer differences between disease groups in the size of stimulus-evoked hemodynamic responses when locomotion co-occurs

As we have previously demonstrated that locomotion can impact hemodynamic responses profoundly^38^, we also assessed stimulus-evoked hemodynamic activity in response to a 2 or 16-s stimulation in trials where locomotion co-occurred at the onset of the stimulus. Where we observed differences between groups to a 2s stimulation presented in the absence of locomotion (Figure 2), when locomotion was also present there were no longer any differences between groups (Figure 4A; SR4), indicating that the vasculature of the atherosclerosis group does have the same capacity to dilate as the other groups, but does not show as large responses to a sensory stimulus when locomotion is not present. Specifically, we observed no difference in the size of the maximum peak of the HbT response (Figure 4B) or the area under the curve of the HbR response (Figure 4C). Since locomotion impacts hemodynamic responses^38^ (Supplementary Figure 6), we plotted the average locomotion traces for trials where locomotion started at the onset of stimulation for each disease group and confirmed that there was no difference in the size of locomotion events between groups (Figure 4D). We conducted the same analysis for the longer duration 16s whisker stimulation (Figure 4E), and again saw no differences between groups in the size of the HbT maximum peak (Figure 4F), HbR area under the curve (Figure 4G) or locomotion events (Figure 4H).

**Fig. 4.**
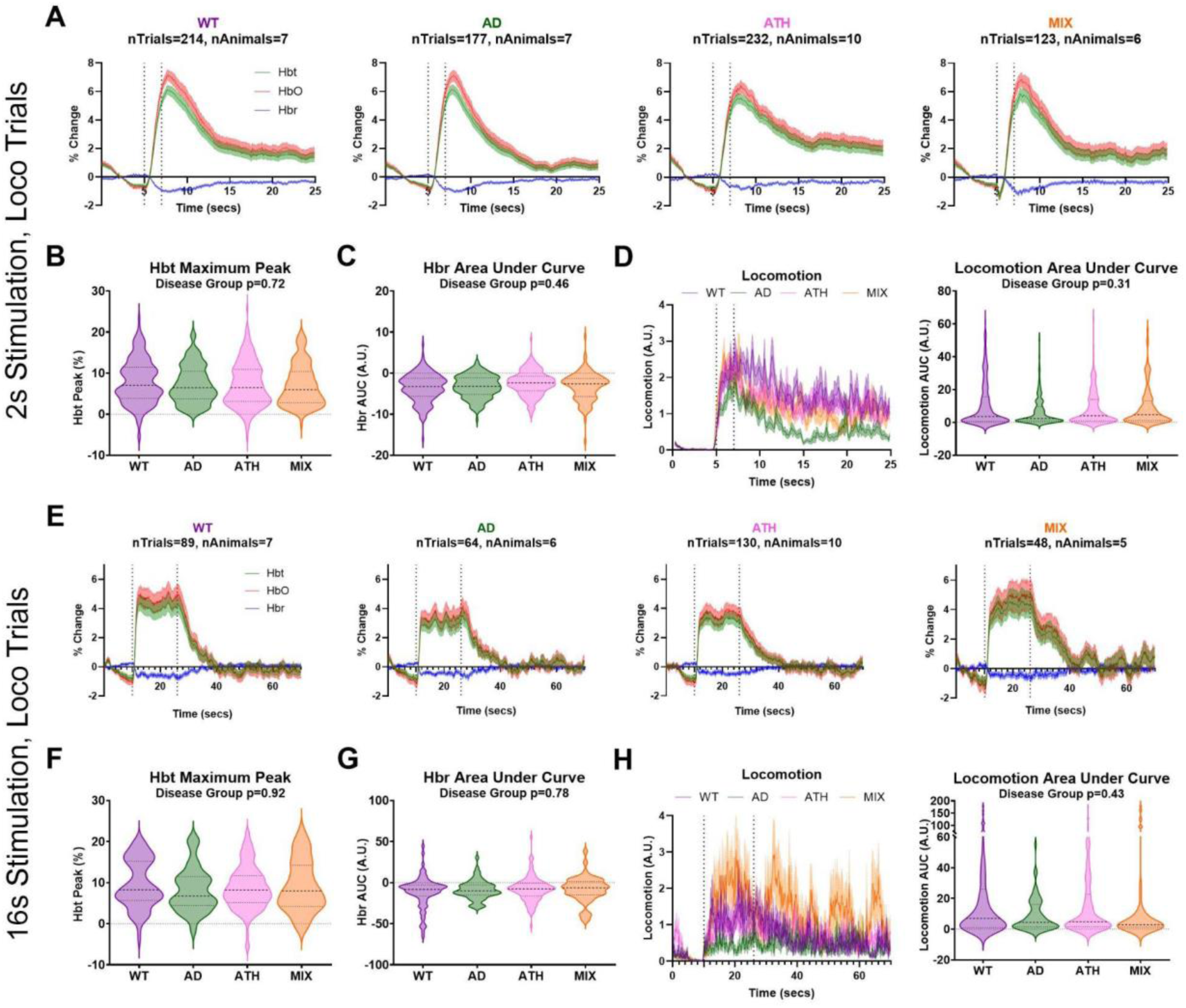
Comparing whisker stimulus-induced responses in trials which contain concurrent locomotion. **A.** Hemodynamic time series from the artery region within the whisker barrels showing total (HbT, green), oxygenated (HbO, red), and deoxygenated (HbR, blue) hemoglobin in response to a 2s mechanical whisker stimulation in trials with concurrent locomotion occurring at the onset of the stimulus period (dotted lines) for wild-type (WT, purple), APP/PS1 (AD, green), atherosclerosis (ATH, pink), and mixed APP/PS1 x atherosclerosis (MIX, orange) mice. There was no difference between disease groups in the size of the **B.** maximum peak of the HbT response (p=0.72), or the **C**. area under the curve of the HbR response (p=0.46) during the stimulation period of these ‘locomotion trials’. **D.** There were no differences in locomotion across disease groups during these ‘locomotion trials’, assessed using the area under the curve of the locomotion response during the stimulation period (right, p=0.31). **E.** Arterial hemodynamic responses were also compared for a 16s mechanical whisker stimulation in trials with concurrent locomotion occurring at the onset of the stimulation period (dotted lines). There was no significant difference in the size of the **F.** HbT (maximum peak, p=0.92) or **G.** HbR (AUC, p=0.78) responses between WT, AD, ATH or MIX mice. **H**. There were also no differences in locomotion during ‘locomotion trials’ for 16s stimulation events (p=0.43). P-values are taken from linear mixed-effects models with disease group inputted as fixed-effect factor, and animal ID as the random effect (lmer package RStudio), and pairwise comparisons (with correction for multiple comparisons) conducted using the Tukey method (emmeans package RStudio). Shaded error bars represent mean +/- SEM. Horizontal lines on violin plots show median and interquartile range. The number of trials and animals included in each group are indicated on the time series graphs.

To confirm our method of locomotion analysis was not skewing our findings regarding hemodynamic responses across disease groups to a 2s whisker stimulation, we also explored another method of locomotion classification where trials were ranked in ascending order from least to most locomotion occurring during the stimulation period- and the bottom and top 20% of locomotion trials were compared (similar to the analysis conducted by Eyre et al., 2022^38^; Supplementary Figure 7). Using this alternative method we replicated our finding that differences were observed between disease groups only in the trials where locomotion was not confounding responses.

### 3.4 Subtle neurovascular deficits are consistent with no effect of disease on recognition memory assessed by the novel object recognition test

We used the novel object recognition test (NOR) to assess recognition memory across groups. Mice were placed into an open field area and could explore two of the same objects (training phase). After a one-hour delay, they were placed back into the same arena, with one of the previous objects replaced with a novel object (testing) (Figure 5a). Mice have an innate preference for novelty, so therefore should want to spend more time with the novel object. Behavioral software was used to assess the time spent with each object (Figure 5b), in order to assess the activity (Figure 5c-d) and preference index (Figure 5e) across groups. Activity was assessed using the distance run (Figure 5c) and velocity of running (Figure 5d) during the NOR training or test session, and we observed no difference between groups indicating that the mice engaged with the task to a similar degree. A higher preference index score indicates a greater preference for the novel object (with scores above 50% indicating performance on the task to distinguish between two objects is above chance). We found no significant differences in the preference index score across disease groups (WT-M: 58.2, SD: 17.1; AD-M: 48.9, SD: 13.5; ATH-M: 61.5, SD: 13.1; MIX-M: 56.9, SD: 17.4; one-way ANOVA F(3,24)=0.88, p=0.47). This suggests that there were no significant differences in the preference that mice had for the novel object – indicating that recognition memory was unchanged in AD, Atherosclerosis, and Mixed disease mice, as compared to WT mice, at 9m of age, with a 1-hour retention interval between training and testing. Although it is worth noting that a 48.9% preference index for the AD mice means that they did not perform above chance overall on the NOR task which could indicate a subtle effect of disease on recognition memory, although our Tukey pairwise comparisons did not show these mice to be performing differently to the other groups (see SR5; AD-WT p=0.71, AD-ATH p=0.39, AD-MIX p=0.79).

**Fig. 5.**
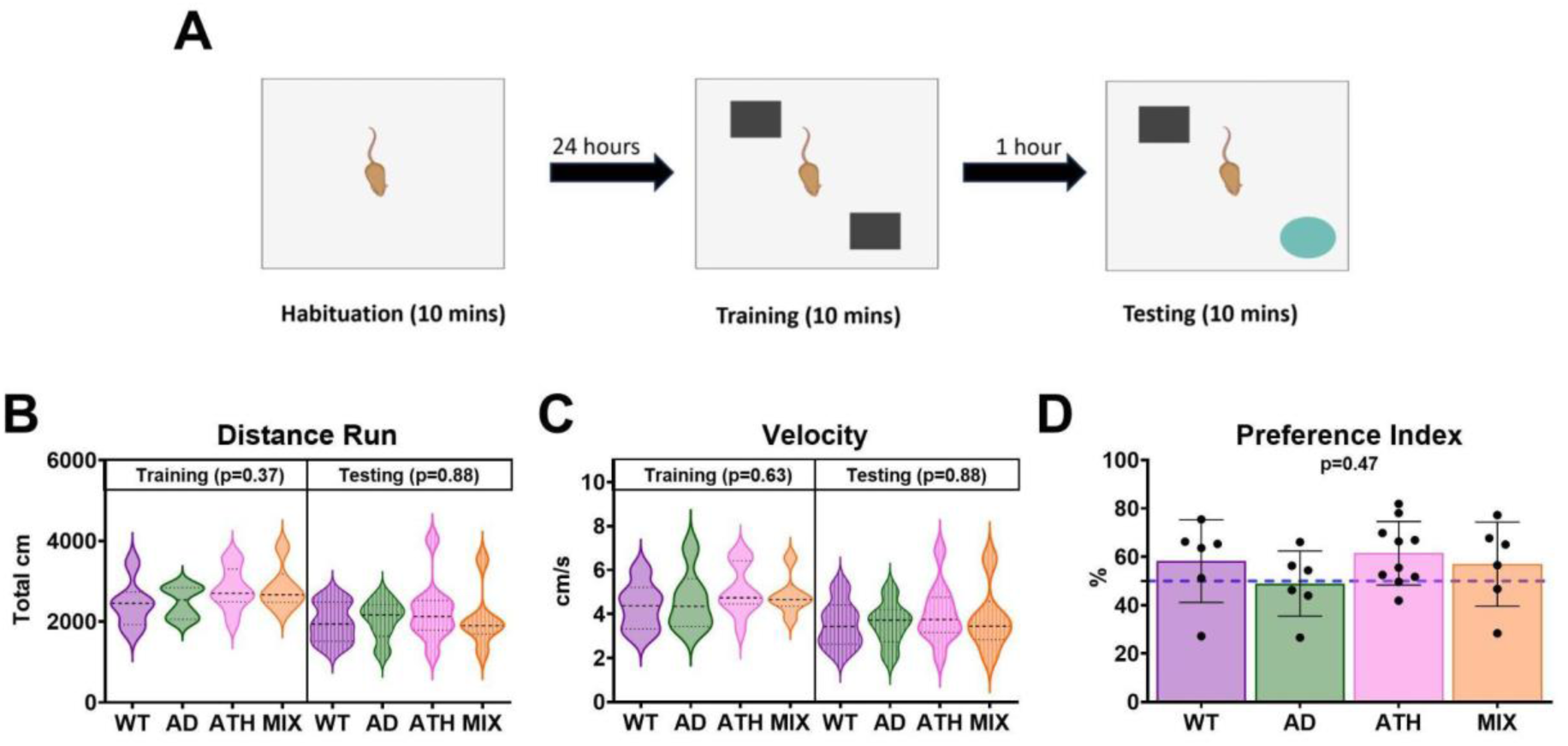
No differences in performance on a novel object recognition task between disease groups. **A.** Mice were placed in a 40x40cm arena and allowed to freely explore for 10-minutes which during the training phase (left) contained two identical objects, and during the testing phase (right) included a novel object to replace either the left or right familiar object from the previous training session (Figure Created using https://BioRender.com). Video recordings were taken of mice in the arena, and they had to explore each object for a minimum of 20 seconds during the training phase to be included in subsequent analysis. We compared the **B.** distance run (cm) and **C.** velocity (cm/s) of each mouse in the training and testing arena to indicate whether mice were engaging with the task similarly between disease groups. We showed no significant difference in the amount of time or speed of travel between groups. **D**. To indicate the mouse’s preference for the novel vs familiar object in the testing arena, the preference index is calculated as: [cumulative duration with novel object / total exploration time (cumulative duration with familiar + novel objects)] x 100. Values over 50% indicate the mouse has distinguished between familiar and novel objects, and spent more time exploring the novel item. The wild-type, atherosclerosis and mixed groups all had values >50% indicating a preference for the novel object, and a one-way ANOVA revealed no overall difference between groups on this task. Because all groups showed a normal distribution, p-values are taken from one-way ANOVAs which group (WT, AD, ATH, MIX) as the independent variable and task performance metric (distance, velocity, preference) as the dependent variable. Shaded error bars represent mean +/- SEM. Horizontal lines on violin plots show median and interquartile range. Individual dots on bar charts represent single mice, and the wild-type group included 6 mice, AD group 6 mice, atherosclerosis group 10 mice, and mixed group 6 mice across all figure panels C-E.

### 3.5 Mixed disease does not enhance atherosclerotic plaque burden or amyloid beta load

As expected, we found atherosclerotic plaques within the aortic arch in atherosclerotic and mixed disease mice (Figure 6a). We assessed whether the presence of amyloid beta within the brain may affect atherosclerosis in the systemic vasculature. We found no significant differences in aortic arch plaque burden between the atherosclerotic and mixed disease mice (Figure 6b, SR6). Amyloid plaque burden was assessed using immunohistochemistry whereby the presence of amyloid was identified using an anti-amyloid antibody with DAB (Figure 6c). We assessed whether the presence of systemic atherosclerosis in addition to AD could impact amyloid coverage in the hippocampus or overlying cortex (Figure 6d; SR6). We found no significant differences in amyloid coverage (calculated as % of total area included in analysis) when comparing AD mice to mixed disease mice (p=0.11) or between brain regions (p=0.99).

**Fig. 6.**
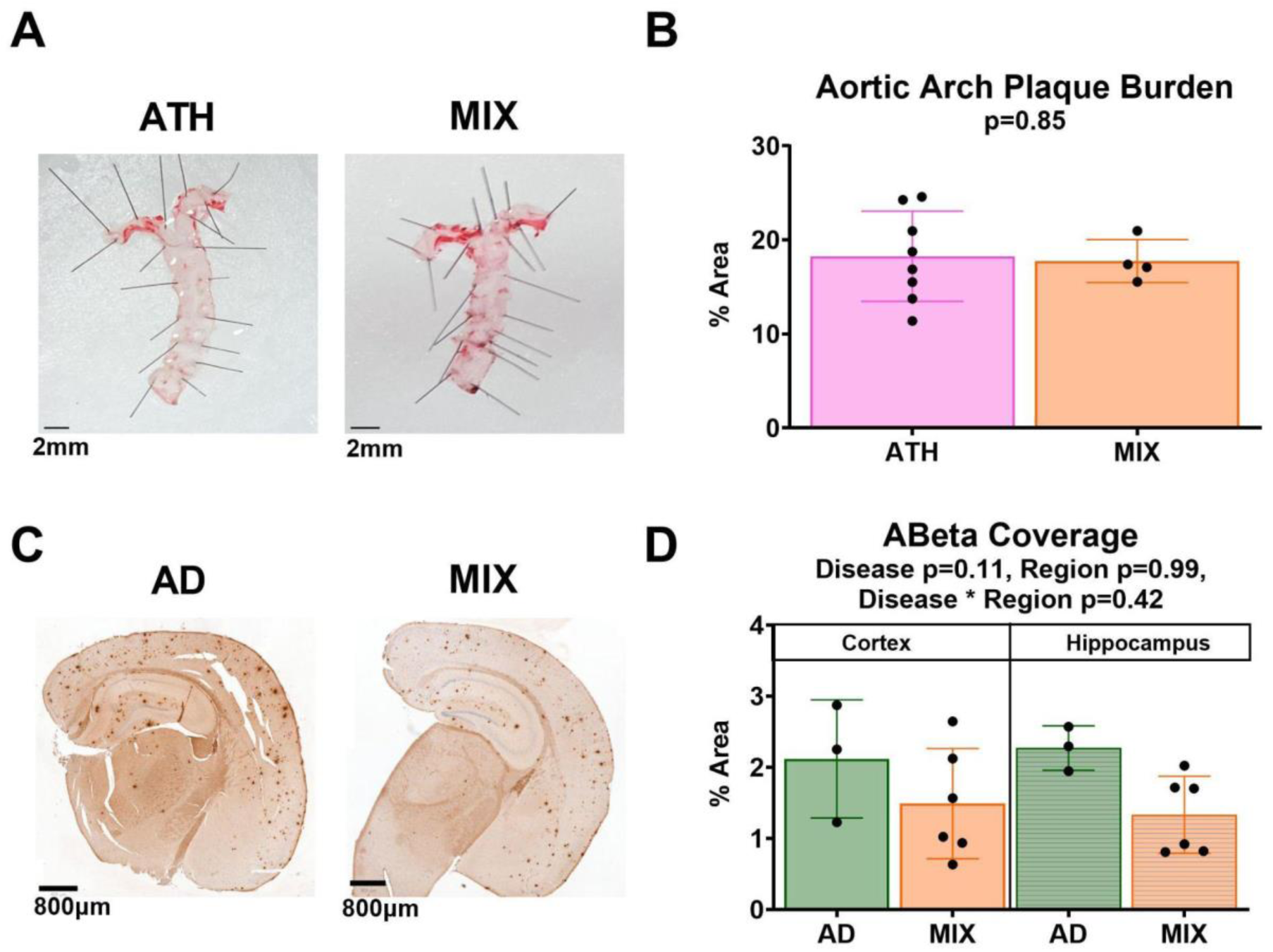
No differences in pathology across disease models. **A.** Representative images of the aorta from an atherosclerotic mouse (left) and a mixed disease mouse (right) from which the percent plaque burden was calculated as the density of plaques found within the area of the aortic arch (top of the image where the aorta branches). The scale bar represents 2mm. **B.** The atherosclerosis (N=8) and mixed disease (N=4) mice showed no differences in plaque burden at the aortic arch (p=0.85). **C.** Representative images of brain slices which include CA1 hippocampus after labelling for amyloid beta in an Alzheimer’s mouse (left) and mixed disease mouse (right). The scale bar represents 800µm. **D.** Regions of interest were taken across the entirety of CA1 for the hippocampus, and for a rectangle overlying CA1 for cortex across all slices and the percentage of amyloid beta was calculated across the total area. There were no significant differences in amyloid coverage across our disease groups (AD N=3, MIX N=6, p=0.11) or brain regions (p=0.99).

## 4 Discussion

In the current study we investigated vascular and cognitive function in Alzheimer’s, atherosclerosis, and mixed disease mouse models. We used 2D-OIS to investigate changes in cortical hemoglobin concentrations to a 2s whisker stimulation, immunohistochemistry to assess hippocampal and cortical amyloid, histology to characterize aortic arch atherosclerotic plaque load, and the NOR task as a test of non-spatial recognition memory.

We observed that in trials in which concurrent locomotion occurred, there was no effect of disease on sensory-evoked hemodynamic responses. However, in support of our previous work, (showing the large effects locomotion can have on hemodynamic responses)^38^ we did find an effect of disease when we controlled for the confounding of locomotion. When only rest trials were considered or when locomotion was ranked during the whisker stimulation, during the least locomotion trials (trials with no locomotion occurring throughout stimulation period, or bottom 20% of trials), we observed that whisker stimulation-evoked peak hemodynamic responses to a 2s-stimulation in the atherosclerosis group were significantly smaller than the wild-type and mixed disease mice. This supported previous work completed by our research group, and therefore extended the findings that peripheral vascular changes, such as atherosclerosis can subtly impact neurovascular function^31^.

Balbi et al. (2015)^42^ previously showed that older wild-type mice (8-12 months) were unable to maintain the size of the initial cerebral blood flow response compared to young mice (6 weeks) when a stimulation was presented up to 10 times. Given we saw disease-related differences in the hemodynamic response for the 2s whisker stimulation (30 trials) but not the 16s whisker stimulation (15 trials), whereby the atherosclerosis mice showed smaller 2s stimulus induced responses, we wanted to ensure this was related to the shorter duration of the stimulus and not the higher number of trials collected. Here we showed that there was no overall difference in the linear relationship between the size of the HbT response and the stimulus trial presentation number across disease groups, meaning the reduced 2s stimulation induced hemodynamic response in atherosclerotic mice was not due to an inability to sustain hemodynamic responses across repeated trials.

We also found preserved recognition memory in all disease groups, as assessed using the NOR test, with a retention interval of 1 hour (although AD mice did not perform above chance on this task, exploring the novel object for only 48.9% of the time spent exploring objects). Finally, our immunohistochemistry experiments to assess amyloid plaque load found no difference in amyloid burden in the mixed disease group compared with the AD only group, as has previously been reported^30,31^, and histology did not reveal differences in plaque burden in the aortic arch of atherosclerosis or mixed disease mice. The similar amyloid plaque burden in the brain of AD and MIX mice was surprising given previous work from our group^31^ showed 3x more plaques in the MIX versus the AD mice, however these differences could be due to the different AD mouse models used (in the previous study the J20 mouse, and here APP/PS1 which has a more severe plaque load), as well as the different analysis methods. Rather than counting the number of plaques as per Shabir et al., we used a semi-automated analysis where the images were binarized and the percent area coverage of amyloid was calculated.

### 4.1 Impact of disease on neurovascular responses

In previous work from our research group, we found that atherosclerosis (but not AD or mixed AD-atherosclerosis) had an effect on evoked-hemodynamic responses in lightly anesthetized mice, whereby we observed smaller whisker-evoked-hemodynamic responses compared to the WT group^31^. In the current study using an awake preparation we also found smaller whisker-evoked-hemodynamic responses in atherosclerotic mice (in a different AD background strain) when mice were not moving and locomotion did not confound stimulation responses. Interestingly, whilst the atherosclerosis mice showed compromised hemodynamic responses, the fact that their hemodynamic responses were preserved to a long duration stimulation (where HbT reached its maximum peak between 3.3-4.6s after stimulus onset, which is outside of the short duration sensory stimulation period) or to a stimulation when locomotion co-occurred indicates that the vasculature itself in the setting of moderate atherosclerosis disease has a preserved capacity to dilate, but that perhaps the signaling mechanisms between the neuron and vessel during the less-sustained short duration stimulus are impacted. It is also interesting that combining AD with atherosclerosis in our hands seemingly ‘rescued’ neurovascular responses. Future work would benefit from an in-depth multi-omics investigation to explore potentially vasodilatory mechanisms impacted across disease groups which could potentially be targeted for benefit in disease.

Whilst it was surprising that we found no effects of AD or mixed AD and atherosclerotic disease on evoked-hemodynamic peak responses in this more ‘severe’ model of AD^32^, previous investigations of hemodynamic responses in APP/PS1 mice which have shown differences in stimulation-evoked hemodynamic responses at 6-months using optical imaging spectroscopy^27^, or reduced arterial dilation to a visual stimulus at 9-12 months^43^, used more invasive surgical procedures and different imaging methods than those used here, and in the second example examined functional hyperemia in a different brain region (visual cortex). Our surgical preparation involving a long recovery period and a thinned window, rather than a craniotomy, was less severe than that used in previous studies. Furthermore, 2D-OIS measurements of global and evoked responses, as used in the current study, are predominantly from the surface vessels of the brain. Other studies using two-photon microscopy in the APP/PS1 mouse model focusing on capillary function have observed a larger number of ‘stalled’ capillaries which, when given a neutrophil antibody, reduced the number of stalls, improving global CBF and short-term memory^44^. Previous work in a different mixed model (LDLR-/-) conducted using two-photon microscopy in APP/PS1 LDLR-/- mice also showed deficits where we did not, but the authors looked at basal changes in individual vessels reporting microvascular morphology changes and reduced tissue oxygenation within single capillaries^27,28^. In the current study we focused on evoked hemodynamic changes in the surface pial vessels of the brain, therefore it could be plausible that there are effects of disease at other locations along the vascular tree.

### 4.2 Impact of disease on cognition

We found no effect of disease on non-spatial recognition memory as assessed by the NOR test, which is in line with the subtle hemodynamic deficits we observed. Whilst our behavioral results are not in line with some previous research in the APP/PS1 model, cognitive measures in mice often produce contradictory results, with some groups finding recognition memory deficits^45,46^ and other groups not^47-49^. It is possible contradictory findings arise from differences in the groups included (e.g. the sex of the mice, the age of the mice, drug treatments or not), the sample sizes used, and the metrics used to assess performance (e.g. location preference, time spent with object, recognition index). Limitations of the current behavioral analysis include the rigid exclusion criteria (2 sessions excluded, and 1 the camera did not record during the task) resulting in some of the groups having a smaller sample. Additionally, the current study only investigated a retention interval of 1 hour, where future work could use both shorter term (e.g., 3 minutes) and longer term (e.g., 24 hour) retention intervals to provide a more extensive understanding of recognition memory in the above disease models.

### 4.3 Importance of accounting for locomotion in awake, behaving experiments

In the current study we observed no effects of disease on evoked-hemodynamic responses when locomotion occurred concurrently with the stimulation. However, a significant effect of disease was observed when no (or the least) locomotion occurred during the whisker stimulation. Previous research has evidenced that locomotion itself can increase hemodynamic responses within the brain^34,38,50,51^. However, our findings may suggest that there are different neurovascular mechanisms controlling locomotion-induced hemodynamic responses compared with whisker-evoked hemodynamic responses, with those underlying the whisker evoked responses being impaired in the atherosclerotic mouse model. Our results show the importance of monitoring locomotion in awake mice, as there may be subtle vascular deficits across diseases, and recording locomotion in order to be able to dissect out the effects of this behavior may allow for the field to fully assess such subtle differences in vascular function across disease groups. Methods of data collection or analysis may need to be adapted to account for locomotion in future investigations, with potentially a larger number of stimulation trials needing to be collected if trials with concurrent locomotion are to be removed (to retain sufficient statistical power to detect differences after data removal), or sophisticated analysis techniques applied to retain the higher number of trials and remove the confounds of locomotion on hemodynamic responses (e.g. using a hemodynamic response function to predict hemodynamic activity from spontaneous behaviors^52^, or applying a kernel analysis or method through which the locomotion-dependent hemodynamic response is subtracted from the hemodynamic response to whisker-stimulation alone).

### 4.4 Conclusion

This study provides a novel understanding of the impact that different neurodegenerative and vascular diseases can have upon the vasculature. For the first time in awake mice we show deficits in the stimulus-evoked hemodynamic responses of atherosclerosis mice when the confound of locomotion is removed. Furthermore, our results emphasize the importance of experimental design when imaging awake mice, whereby stimulus duration should be considered and locomotion monitored.

## 6 Code, Data, and Materials Availability

- The data presented in this article are publicly available on ORDA at [DOI link to be added prior to publication]. The individual hemodynamic and locomotion traces with corresponding labels (vessel group, disease group, imaging session, trial number) are available as MATLAB files, and the time series parameters used for the statistical calculations as a .csv file.
- The custom code used for data analysis is available as a Github repository (kirashaw1/CharacterizingVascFunction_ADATH), which has been published via Zenodo (https://doi.org/10.5281/zenodo.14238624).
- The outputs of all statistical analyses are reported in statistical tables SR1-6 in Appendix B.

## Acknowledgements

We thank Rachel Sandy at the University of Sheffield for expertise regarding IV injections as well as the whole BSU team for their husbandry expertise. CH was funded by a Sir Henry Dale Fellowship jointly funded by the Wellcome Trust and the Royal Society. This research was funded in whole, or in part, by the Wellcome Trust [Grant number 105586/Z/14/Z]. The research was also funded by a Batelle-Jeff Wadsworth PhD Studentship (BE) and a UKRI MRC grant [Grant number MR/X003418/1]. For the purpose of open access, the author has applied a Creative Commons Attribution (CC BY) licence to any Author Accepted Manuscript version arising.

## Author Biographies

**Beth Eyre** is a research fellow at Massachusetts General Hospital/Harvard Medical School. She received her BSc degree (Psychology, *International*) from the University of Leeds in 2017, her MSc in Cognitive Neuroscience and human Neuroimaging from the University of Sheffield in 2020 and also her PhD in 2023. She is an author on 4 journal papers and her current research interests include cerebral amyloid angiopathy, brain clearance and neurovascular coupling.

**Kira Shaw** is a postdoctoral research associate at the University of Sheffield. She received her BS degree in Psychology from the University of Leeds in 2011, her MS Psychology degree from the University of Manchester in 2012, and her PhD in Neuroscience from the University of Sheffield in 2017. She is an author on 10 journal papers, and her current research interests include studying neurovascular coupling and the impact perturbations in energy supply may place on brain function.

**Jason Berwick** is a Reader in Neurophysiology and Neuroimaging at the University of Sheffield who specializes in the use of in vivo multimodal brain imaging and electrophysiological methodologies to understand the mechanisms of neurovascular coupling in health and disease. He has published over 60 research papers on the topic of NVC.

**Clare Howarth** is a Senior Lecturer at the University of Sheffield. She received her MSci degree in Physics with a year in Europe from Imperial College London in 2003, and her PhD in Neuroscience from University College London in 2008. She is an author on 29 papers, and her current research interests focus on neurovascular coupling in health and disease.

Biographies and photographs for the other authors are not available.

## Caption List

**S Fig. 1** Female mice show significantly smaller HbR washout in the artery region.

**S Fig. 2** Female mice show smaller hemodynamic responses in the parenchymal region.

**S Fig. 3** Comparison of sex-dependent performance on the novel object recognition task.

**S Fig. 4** 2s stimulus responses without the confounds of locomotion across the different vascular compartments.

**S Fig. 5** 16s stimulus responses without the confounds of locomotion across the different vascular compartments.

**S Fig. 6** Correlating locomotion and hemodynamic responses.

**S Fig. 7** Assessing the impact of locomotion on 2s-stimulus induced hemodynamic responses across ranked trials.

