## Supplementary data for "Characterizing vascular function in mouse models of Alzheimer’s disease, atherosclerosis, and mixed Alzheimer’s and atherosclerosis"

### Appendix A: Supplemental Material

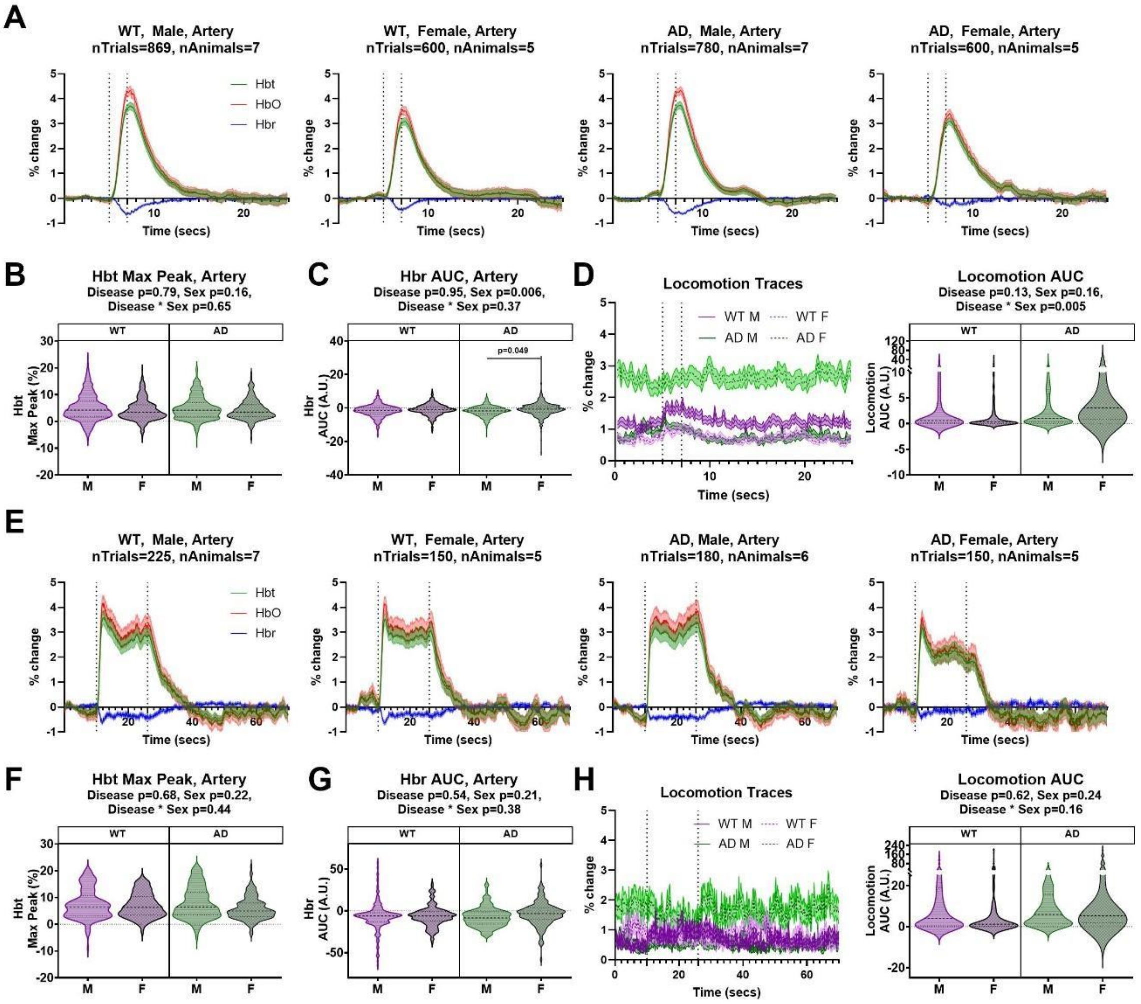


**S Fig. 1** Female mice show significantly smaller HbR washout in the artery region.

**A.** Hemodynamic time series are visualised for WT male (left), WT female (centre left), AD male (centre right) and AD female (right) mice in response to a 2s mechanical whisker stimulation. The size of hemodynamic responses by disease group (WT/AD) or sex (M/F) in the artery ROI were assessed using the **B.** Hbt maximum peak, and **C.** Hbr area under the curve. A linear mixed model analysis revealed the HbR AUC showed a significant impact of sex (p=0.006), with females showing smaller responses than males. **D.** The size of the locomotion events was found to be significantly different, with female AD mice running more than the other groups (disease * sex interaction p=0.005), although this would not impact the differences observed in the hemodynamics as if anything would increase responses in this group. **E.** 16s hemodynamic time series were also visualized across groups, and the size of **F.** Hbt, **G.** Hbr, and **H.** locomotion responses compared. There was found to be no significant effect of disease group or sex on hemodynamic responses in the linear mixed model, although when the distribution of the individual datasets were compared directly using a Kruskall-Wallis test there was a significant difference in the size of the Hbt maximum peak (p=0.013) and Hbr AUC (p=0.002), with the AD female group showing the smallest responses.

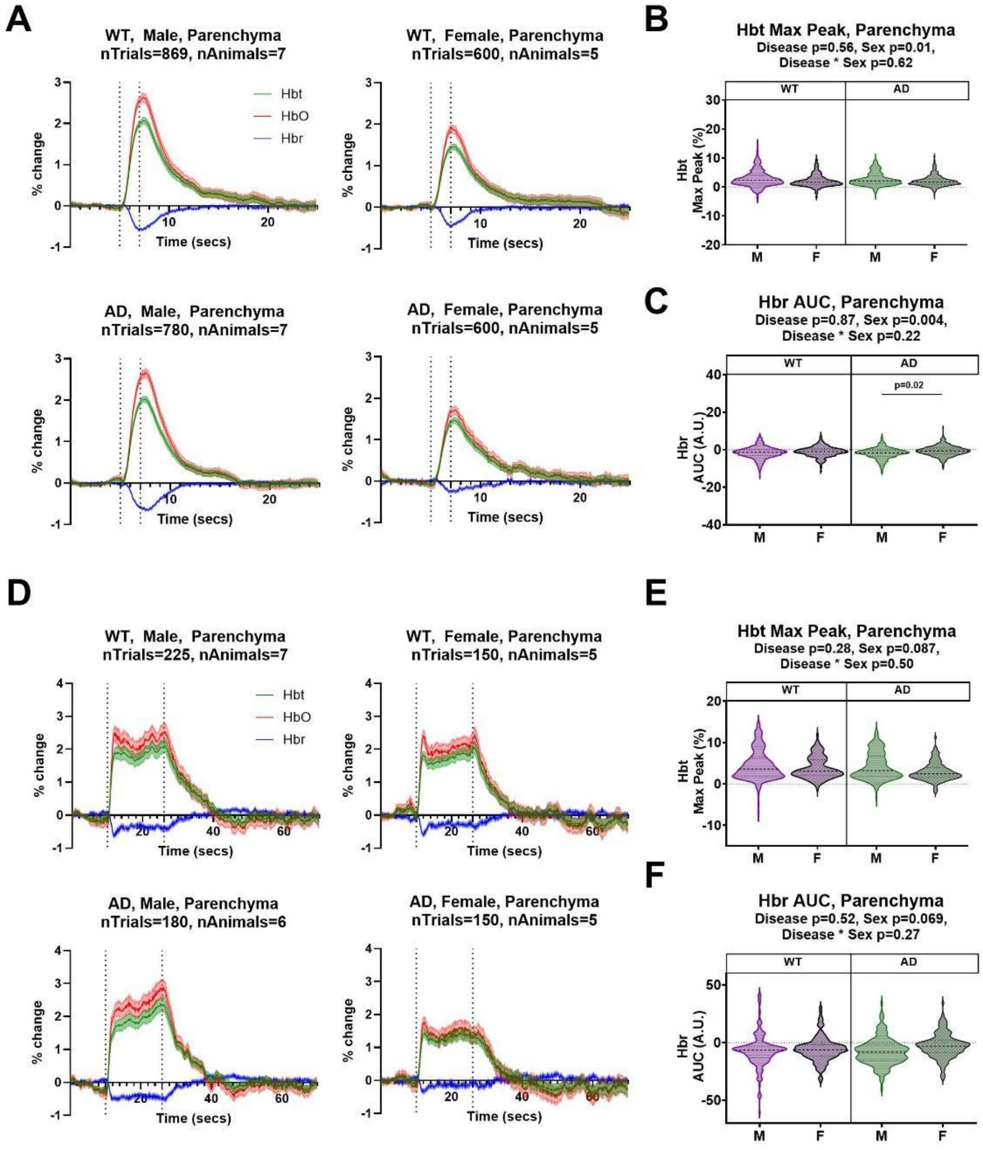


**S Fig. 2** Female mice show smaller hemodynamic responses in the parenchymal region.

**A.** Hemodynamic traces to a 2s whisker stimulation were also visualised for the parenchymal region (which is reflective of blood volume changes from the capillary bed) for WT male (top left), WT female (top right). AD male (bottom left), and AD female (bottom right) mice. The size of hemodynamic responses by disease group (WT/AD) or sex (M/F) in the parenchyma ROI were assessed using linear mixed models for the **B.** HbT maximum peak, and **C.** HbR area under the curve. There was a significant impact of sex on the magnitude of both the HbT (p=0.01) and HbR (p=0.004) responses, with females showing smaller changes from baseline to the whisker stimulation than males. **D.** The hemodynamic responses to a 16s whisker stimulation in the parenchymal region revealed trend level (p<0.1) sex differences in the size of the **E.** HbT maximum peak (sex p=0.087) and **F.** HbR area under the curve (p=0.069), with females again showing smaller responses than males overall.

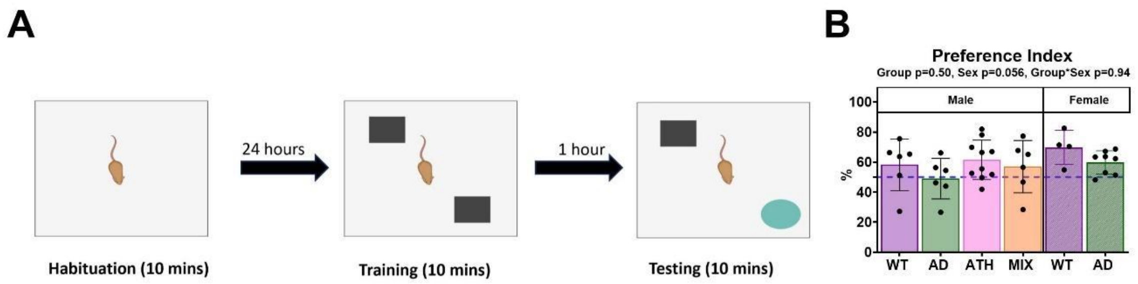


**S Fig. 3** Comparison of sex-dependent performance on the novel object recognition task.

**A.** Male and female wild-type (WT) and APP/PS1 Alzheimer's disease (AD) mice, and male atherosclerosis (ATH) and mixed atherosclerosis/AD (MIX) were exposed to a novel object recognition task consisting of a 10-minute training phase, before the 10-minute testing protocol occurred where a novel object was introduced (Figure created in https://BioRender.com). **B.** The preference index was calculated during the testing phase for all eligible mice, with scores >50% indicating more time spent with the novel object than the familiar object. There was a trend level difference in performance, with female mice performing better on the task than male mice overall due to a higher preference for novelty (p=0.056).

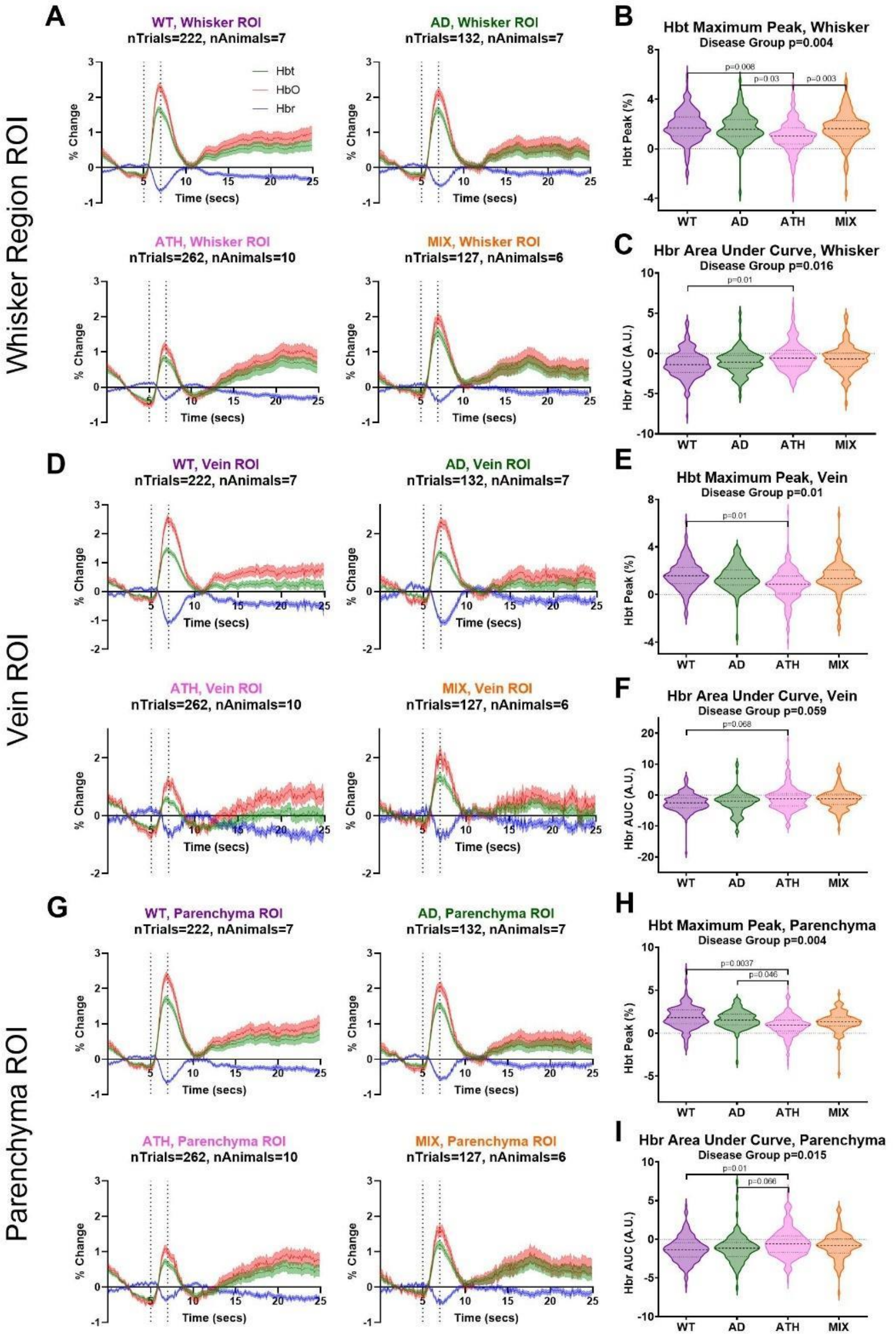


**S Fig. 4** 2s stimulus responses without the confounds of locomotion across the different vascular compartments. Hemodynamic time series showing total (HbT, green), oxygenated (HbO, red), and deoxygenated (HbR, blue) hemoglobin in response to a 2s mechanical whisker stimulation in trials with no concurrent locomotion occurring between the 4 seconds either side of the stimulus period (dotted lines) for wild-type (WT, purple), APP/PS1 (AD, green), atherosclerosis (ATH, pink), and mixed APP/PS1 x atherosclerosis (MIX, orange) mice were also shown for the whisker (top, **A.**), vein (middle, **D.**) and parenchyma (bottom, **G.**) regions of interest. Findings were consistent with those observed in Figure 1, and showed a significant impact of disease group on the size of HbT and HbR responses which was driven by smaller responses in the atherosclerosis group (**B-C, E-F, H-I**).

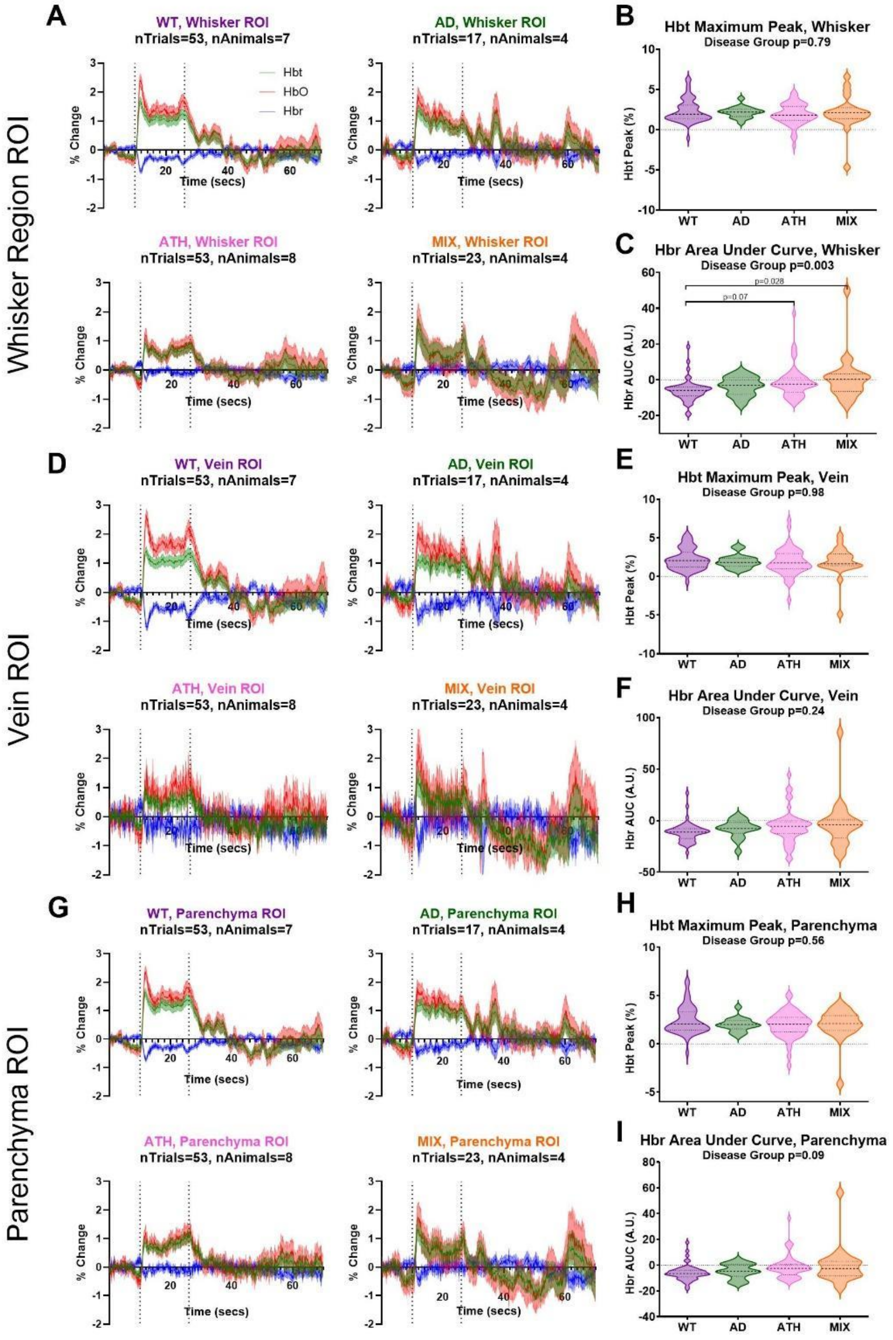


**S Fig. 5** 16s stimulus responses without the confounds of locomotion across the different vascular compartments.

Hemodynamic time series showing total (HbT, green), oxygenated (HbO, red), and deoxygenated (HbR, blue) hemoglobin in response to a 16s mechanical whisker stimulation in trials with no concurrent locomotion occurring between the 4 seconds either side of the stimulus period (dotted lines) for wild-type (WT, purple), APP/PS1 (AD, green), atherosclerosis (ATH, pink), and mixed APP/PS1 x atherosclerosis (MIX, orange) mice were also shown for the artery (top, **A**.), vein (middle, **D**.) and parenchyma (bottom, **G.**) regions of interest (which were taken from within the larger whisker barrel ROI). Findings were consistent with those observed in Figure 1, and showed no significant impact of disease group on the size of HbT and HbR responses (**B-C, E-F, H-I**).

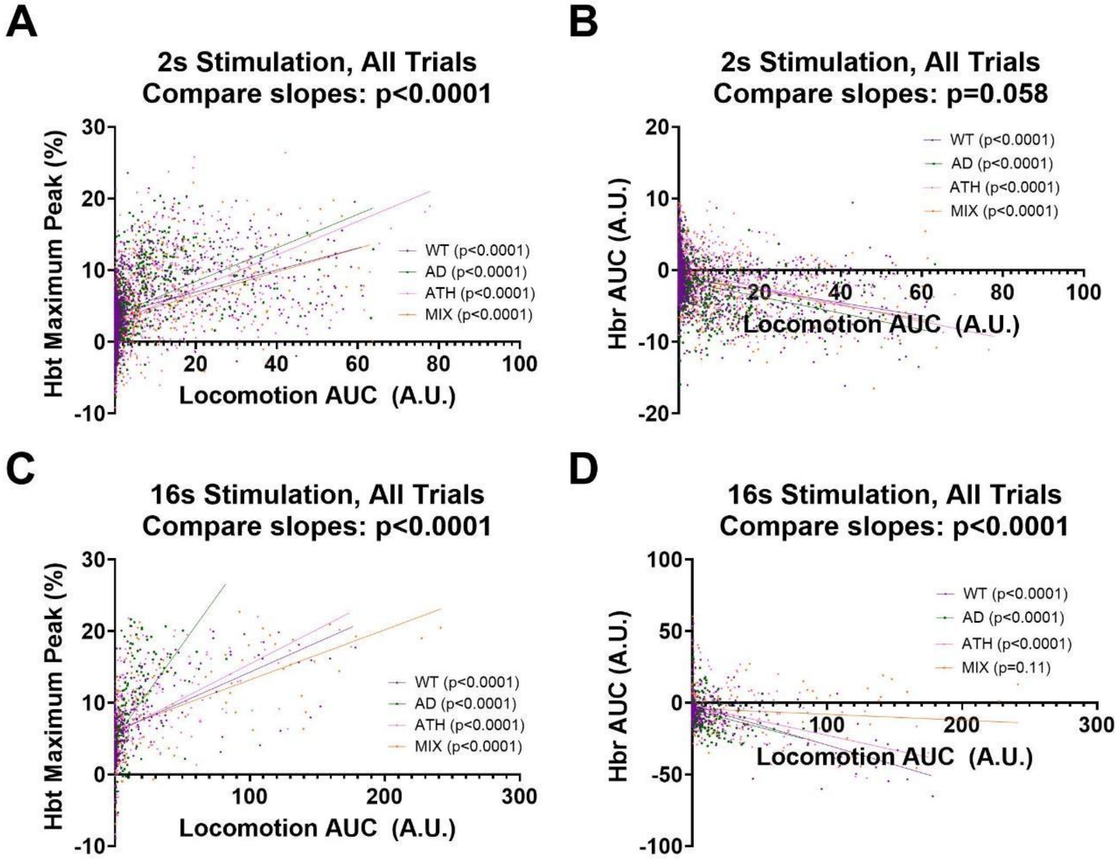


**S Fig. 6** Correlating locomotion and hemodynamic response

The impact of locomotion on hemodynamic responses was visualised across all trials using scatterplots for the 2s-stimulation induced **A.** HbT maximum peak and **B.** HbR area under the curve, and the 16s-stimulation induced **C.** HbT maximum peak and **D.** HbR area under the curve. Across all scatterplots, Pearson's correlations were conducted within each disease group to assess whether locomotion and hemodynamic responses correlated (p-values displayed in legend), and simple linear regressions conducted to look for an overall effect of disease (comparing slopes between groups, p-values displayed in the title). There was a consistent effect of locomotion on hemodynamic responses across all groups, with locomotion increasing the size of responses.

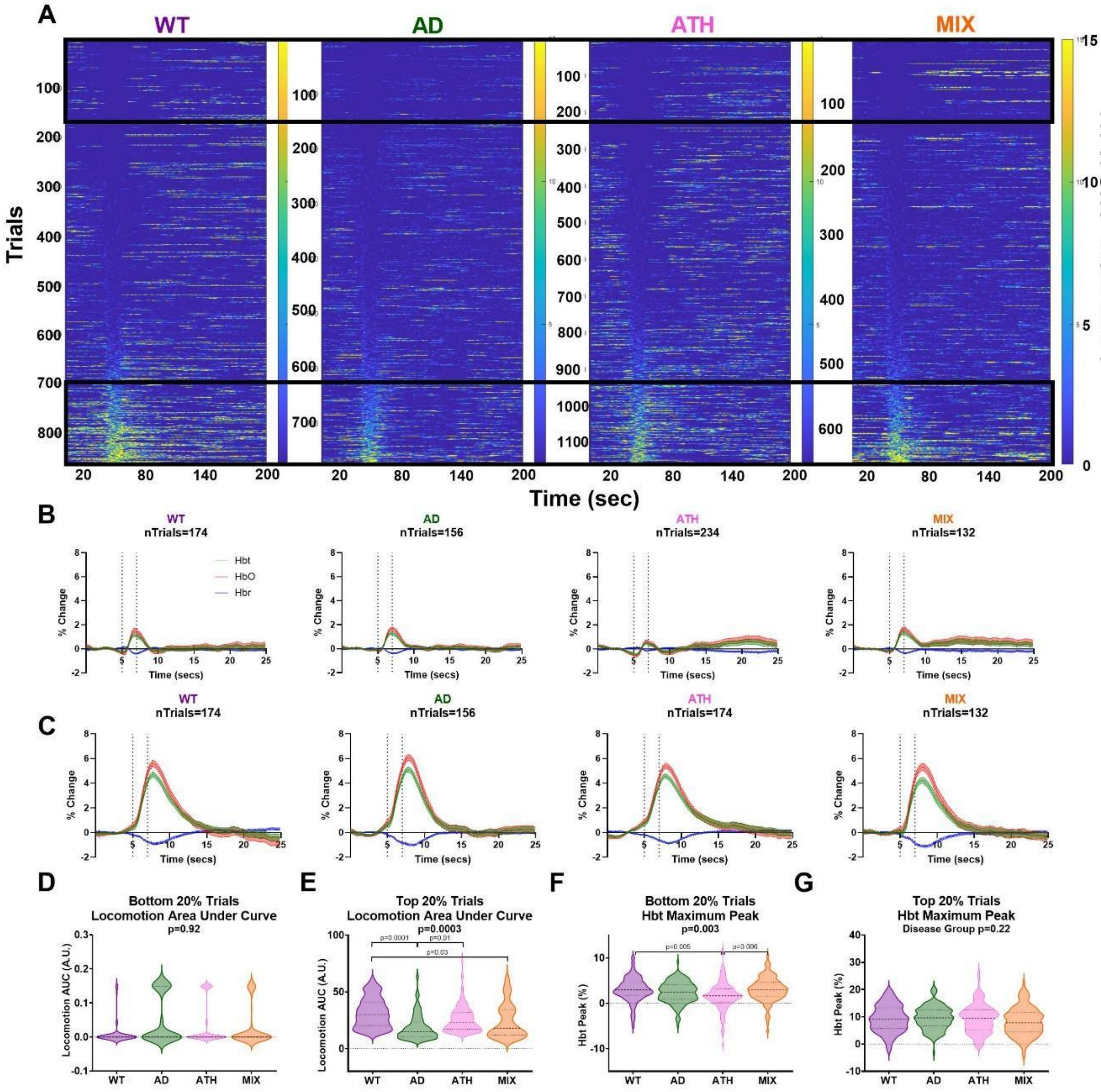


**S Fig. 7** Assessing the impact of locomotion on 2s-stimulus induced hemodynamic responses across ranked trials.

An alternative method of assessing the impact of locomotion on hemodynamic responses was explored (vs Figures 2 and 4), as **A.** individual trials were ranked in ascending order from least to most locomotion during the stimulation period for WT (left, purple), AD (centre left, green), ATH (centre right, pink) and MIX (right, orange) mice. The hemodynamic responses to a 2s-whisker stimulation where then visualised for the **B.** bottom and **C.** top 20% of ranked trials across all disease groups. Across disease groups the size of locomotion responses was **D.** not different in the bottom 20% of trials (as there was no running for any of the groups), **E.** but was significantly different in the top 20% of trials (p=0.0003), where WT animals ran more than the other groups. However, using this alternative method of understanding the impact of locomotion on hemodynamic responses, findings were consistent to those observed previously, with **F.** a significant effect of disease observed in the size of the HbT response when no locomotion was present (p=0.003) due to atherosclerosis mice showing smaller responses, and **G.** no effect of disease on HbT responses impacted by locomotion (top 20% of trials).

#### Appendix B: Statistical Reports

**Statistics Reports (SR1-SR6):** The following tables report the mean, standard deviation, type of statistical test, test statistics, degrees of freedom and p-value for each statistical test reported in the main manuscript.

**SR1: Figure 2: Comparing stimulation-induced responses without the confounds of locomotion.** The time series metrics across disease groups (4 levels: WT, AD, ATH, MIX) for the stimulus-induced responses in the artery ROI from the whisker barrel region where compared in figures 2b-d & f-h using a linear mixed model (lmer package RStudio) with disease group as the independent variable, the dependent variable as HbT maximum peak (b,f), HbR area under the curve (c,g) or locomotion area under the curve (d,h), and animal ID as the random factor (to account for variations between groups being driven by a single outlier animal). To further investigate where the specific significant differences were pairwise comparisons (with correction for multiple comparisons) were conducted using the Tukey method (emmeans function RStudio).

| **Figure** | **Mean** | **SD** | **Test** | **Test Statistic** | **Mean Square** | **Degrees of freedom** | **P Value** | **Post-hoc comparisons:** |
|---|---|---|---|---|---|---|---|---|
| 2b (HbT max peak, 5-10s) | WT: 3.256, AD: 2.985, ATH: 1.905, MIX: 3.310 | WT: 2.441, AD: 2.195, ATH: 2.597, MIX: 2.590 | Linear mixed model (fixed factor: disease group) | F=4.97 | 28.689 | 3 | 0.0081 | AD-ATH p=0.1195, AD-MIX p=0.8210, AD-WT p=0.8886, ATH-MIX p=0.0267, ATH-WT p=0.0179, MIX-WT p=0.9945 |
| 2c (HbR AUC, 5-10s) | WT: -1.261, AD: -1.275, ATH: -0.3855, MIX: -0.6113 | WT: 1.899, AD: 2.015, ATH: 2.163, MIX: 1.942 | Linear mixed model (fixed factor: disease group) | F=3.746 | 14.593 | 3 | 0.0303 | AD-ATH p=0.0547, AD-MIX p=0.5851, AD-WT p=1.00, ATH-MIX p=0.7145, ATH-WT p=0.0416, MIX-WT p=0.5536 |
| 2d (Locomotion AUC, 5-10s) | WT: 0.08118, AD: 0.08145, ATH: 0.09893, MIX: 0.06603 | WT: 0.1282, AD: 0.1213, ATH: 0.1416, MIX: 0.1163 | Linear mixed model (fixed factor: disease group) | F=1.2738 | 0.021166 | 3 | 0.3115 | AD-ATH p=0.6468, AD-MIX p=0.9403, AD-WT p=0.9999, ATH-MIX p=0.3360, ATH-WT p=0.5210, MIX-WT p=0.9465 |

| | | | | | | | | |
|---|---|---|---|---|---|---|---|---|
| 2f (HbT max peak, 10-30s) | WT: 4.563, AD: 3.540, ATH: 3.329, MIX: 4.328 | WT: 3.033, AD: 1.288, ATH: 2.594, MIX: 3.768 | Linear mixed model (fixed factor: disease group) | F=0.6258 | 4.1957 | 3 | 0.6058 | AD-ATH p=0.9967, AD-MIX p=0.9226, AD-WT p=0.8920, ATH-MIX p=0.7655, ATH-WT p=0.6445, MIX-WT p=1.00 |
| 2g (HbR AUC, 10-30s) | WT: -5.667, AD: -4.17, ATH: -1.337, MIX: 1.375 | WT: 6.815, AD: 5.511, ATH: 10.07, MIX: 14.94 | Linear mixed model (fixed factor: disease group) | F=3.3007 | 302.19 | 3 | 0.05927 | AD-ATH p=0.7404, AD-MIX p=0.3326, AD-WT p=0.9535, ATH-MIX p=0.7144, ATH-WT p=0.1983, MIX-WT p=0.0632 |
| 2h (Locomotion AUC, 10-30s) | WT: 0.1447, AD: 0.2929, ATH: 0.1634, MIX: 0.1032 | WT: 0.1888, AD: 0.2381, ATH: 0.1790, MIX: 0.1405 | Linear mixed model (fixed factor: disease group) | F=1.9185 | 0.05847 | 3 | 0.1978 | AD-ATH p=0.3405, AD-MIX p=0.1363, AD-WT p=0.2522, ATH-MIX p=0.7748, ATH-WT p=0.9905, MIX-WT p=0.8950 |

**SR2: Figure 3: Differences across disease models in the size of 2s-stimulation responses during rest are not linked to the high number of trials.** The impact of trial number on the size of HbT responses (HbT maximum peak or HbR area under the curve) was assessed using a linear regression across disease groups. Generally for each mouse group (WT, AD, ATH, MIX) there was no relationship between the size of the HbT response and the trial number (see is the slope significantly non-zero), except for the wild-type animals in the size of the HbT maximum peak which increased as trial number increased (positive correlation), and for the MIX animals in the size of the HbR AUC which increased (larger decrease) as the trial number increased (negative correlation). There was no overall significant difference between the disease groups on the relationship between locomotion and the HbT responses (see are the slopes equal), but the intercepts were significantly different between groups (see are the intercepts equal) as the HbT maximum peak values were generally smaller for ATH mice (intercepts y-axis at lower value), and HbR AUC values generally smaller in the earlier trials for the MIX mice (intercepts y-axis closer to 0).

| **Figure** | **Test** | **Is the slope significantly non-zero?** | **Are the slopes equal?** | **Are the intercepts equal?** |
|---|---|---|---|---|

| | | | | |
|---|---|---|---|---|
| 3a HbT maximum peak/ trial number | Linear Regression | WT: F(1,220)=5.160, p=0.0241<br><br>AD: F(1,130)=0.6367, p=0.4263<br><br>ATH: F(1,260)=0.02847, p=0.8661<br><br>MIX: F(1,125)=2.466, p=0.1188 | F(3,735)=1.799, p=0.1460 | F(3,738)=15.20, p<0.0001 |
| 3d HbR AUC / trial number | Linear Regression | WT: F(1,220)=2.391, p=0.1235<br><br>AD: F(1,130)=0.07866, p=0.7796<br><br>ATH: F(1,260)=0.6943, p=0.4055<br><br>MIX: F(1,125)=7.138, p=0.0086 | F(3,735)=1.130, p=0.3360 | F(3,738)=10.00, p<0.0001 |

**SR3: Figure 3: Differences across disease models in the size of 2s-stimulation responses during rest are not linked to the high number of trials.** The impact of disease group (WT, AD, ATH, MIX) and trial number on the size of HbT and HbR responses was assessed by categorizing data as belonging to early (trials 1-5) or late (trials 25-30) trials, and comparing the size of the responses (3b. maximum peak for HbT, and 3e. area under the curve for HbR). A linear mixed model (lmer package RStudio) was conducted with disease group and trial group (early or late) as the independent variables, and the dependent variable as HbT maximum peak (b) or HbR area under the curve (e), and animal ID as the random factor (to account for variations between groups being driven by a single outlier animal). To further investigate where the specific trend-level differences were pairwise comparisons (with correction for multiple comparisons) were conducted using the Tukey method (emmeans function RStudio). Through these linear mixed models we observed no significant impact of trial number on the size of hemodynamic responses.

| **Figure** | **Mean** | **SD** | **Test** | **Test Statistic** | **Mean Square** | **Degrees of freedo m** | **P Value** | **Post-hoc comparisons:** |
|---|---|---|---|---|---|---|---|---|

| | | | | | | | | |
|---|---|---|---|---|---|---|---|---|
| 3b (HbT maximum peak) | WT early: 2.022, AD early: 3.266, ATH early: 2.553, MIX early: 3.264, WT late: 3.801, AD late: 2.969, ATH late: 2.412, MIX late: 3.835 | WT early: 2.733, AD early: 1.778, ATH early: 3.098, MIX early: 2.242, WT late: 2.534, AD late: 1.720, ATH late: 1.780, MIX late: 2.835 | Linear mixed model (fixed factors: disease group* trial number ) | Disease group: 0.9297, trial group: 1.6510, Disease group * trial group: 2.3865 | Disease group: 4.9543, trial group: 8.7977, Disease group * trial group: 12.7167 | Disease group: 3, trial group: 1, Disease group * trial group: 3 | Disease group: p=0.4416 , trial group: p=0.200, Disease group * trial group: p=0.0696 | AD Early - ATH Early p=0.9928<br>AD Early - MIX Early p=1.0000<br>AD Early - WT Early p=0.8890<br>AD Early - AD Late p=0.9999<br>AD Early - ATH Late p=0.9585<br>AD Early - MIX p=0.9947<br>AD Early - WT p=0.9940<br>ATH Early - MIX p=0.9943<br>ATH Early - WT Early p=0.9966<br>ATH Early - AD Late p=0.9999<br>ATH Early - ATH Late p=0.9999<br>ATH Early - MIX Late p=0.6818<br>ATH Early - WT Late p=0.5760<br>MIX Early - WT Early p=0.9238<br>MIX Early - AD Late p=0.9999<br>MIX Early - ATH Late p=0.9741<br>MIX Early - MIX Late p=0.9988 |

| | | | | | | | | |
|---|---|---|---|---|---|---|---|---|
| | | | | | | | | MIX Early - WT Late p=0.9997<br><br>WT Early - AD Late p=0.9691<br><br>WT Early - ATH Late p=0.9998<br><br>WT Early - MIX Late p=0.4089<br><br>WT Early - WT Late p=0.1077<br><br>AD Late - ATH Late p=0.9955<br><br>AD Late - MIX Late p=0.9337<br><br>AD Late - WT Late p=0.9122<br><br>ATH Late - MIX Late p=0.4816<br><br>ATH Late - WT Late p=0.3557<br><br>MIX Late - WT Late p=1.0000 |

| | | | | | | | | |
|---|---|---|---|---|---|---|---|---|
| 3e (HbR AUC) | WT early: -1.026, AD early: -0.7663, ATH early: -0.3487, MIX early: -0.3744, WT late: -1.471, AD late: -1.144, ATH late: -0.8161, MIX late: -1.258 | WT early: 2.227, AD early: 1.337, ATH early: 2.360, MIX early: 1.453, WT late: 1.895, AD late: 1.325, ATH late: 2.022, MIX late: 2.259 | Linear mixed model (fixed factors: disease group* trial number) | Disease group: 1.1221, trial group: 3.4293, Disease group * trial group: 0.1470 | Disease group: 4.199, trial group: 12.8354, Disease group * trial group: 0.5502 | Disease group: 3, trial group: 1, Disease group * trial group: 3 | Disease group: p=0.36178, trial group: p=0.06521, Disease group * trial group: p=0.93155 | AD Early - ATH Early -0.3913 0.592 87.2 -0.661 0.9978<br><br>AD Early - MIX Early p=0.9999<br><br>AD Early - WT Early p=1.0000<br><br>AD Early - AD Late p=0.9979<br><br>AD Early - ATH Late p=1.0000<br><br>AD Early - MIX Late p=0.9939<br><br>AD Early - WT Late p=0.8943<br><br>ATH Early - MIX Early p=1.0000<br><br>ATH Early - WT Early p=0.9628<br><br>ATH Early - AD Late p=0.8357<br><br>ATH Early - ATH Late p=0.9942<br><br>ATH Early - MIX Late p=0.7284<br><br>ATH Early - WT Late p=0.2564<br><br>MIX Early - WT Early |

| | | | | | | | | |
|---|---|---|---|---|---|---|---|---|
| | | | | | | | | p=0.9972<br><br>MIX Early - AD Late p=0.9833<br><br>MIX Early - ATH Late p=1.0000<br><br>MIX Early - MIX Late p=0.9381<br><br>MIX Early - WT Late p=0.8132<br><br>WT Early - AD Late p=1.0000<br><br>WT Early - ATH Late p=0.9995<br><br>WT Early - MIX Late p=0.9998<br><br>WT Early - WT Late p=0.9648<br><br>AD Late - ATH Late p=0.9877<br><br>AD Late - MIX Late p=1.0000<br><br>AD Late - WT Late p=0.9962<br><br>ATH Late - MIX Late p=0.9587<br><br>ATH Late - WT Late |

| | | | | | | | | |
|---|---|---|---|---|---|---|---|---|
| | | | | | | | | p=0.6138<br><br>MIX Late - WT Late<br>p=0.9996 |

**SR4: Figure 4: Comparing stimulation-induced responses which contain concurrent locomotion.** The time series metrics across disease groups (4 levels: WT, AD, ATH, MIX) for the stimulus-induced responses in the artery ROI from the whisker barrel region during a 16s stimulation where compared in figures 4b-d & f-h using a linear mixed model (lmer package RStudio) with disease group as the independent variable, the dependent variable as HbT maximum peak (b,f), HbR area under the curve (c,g) or locomotion area under the curve (d,h), and animal ID as the random factor (to account for variations between groups being driven by a single outlier animal). To further investigate where the specific significant differences were pairwise comparisons (with correction for multiple comparisons) were conducted using the Tukey method (emmeans function RStudio).

| **Figure** | **Mean** | **SD** | **Test** | **Test Statistic** | **Mean Square** | **Degrees of freedom** | **P Value** | **Post-hoc comparisons:** |
|---|---|---|---|---|---|---|---|---|
| 4b (HbT max peak, 5-10s) | WT: 7.917, AD: 7.257, ATH: 7.196, MIX: 7.058 | WT: 5.318, AD: 4.536, ATH: 5.387, MIX: 5.006 | Linear mixed model (fixed factor: disease group) | F=0.4526 | 10.685 | 3 | 0.7177 | AD-ATH p=0.9984, AD-MIX p=0.9995, AD-WT p=0.7778, ATH-MIX p=0.9931, ATH-WT p=0.8236, MIX-WT p=0.7538 |
| 4c (HbR AUC, 5-10s) | WT: -3.520, AD: -3.314, ATH: -2.465, MIX: -3.313 | WT: 3.373, AD: 2.736, ATH: 2.947, MIX: 3.411 | Linear mixed model (fixed factor: disease group) | F=0.8817 | 7.5197 | 3 | 0.4634 | AD-ATH p=0.6520, AD-MIX p=0.9988, AD-WT p=0.9911, ATH-MIX p=0.7953, ATH-WT p=0.4543, MIX-WT p=0.9742 |
| 4d (Locomotion AUC, 5-10s) | WT: 10.55, AD: 6.574, ATH: 8.593, MIX: 9.151 | WT: 13.67, AD: 8.509, ATH: 10.17, MIX: 11.57 | Linear mixed model (fixed factor: disease group) | F=1.248 | 145.26 | 3 | 0.3129 | AD-ATH p=0.8100, AD-MIX p=0.5737, AD-WT p=0.2783, ATH-MIX p=0.9413, ATH-WT p=0.6997, MIX-WT p=0.9790 |

| | | | | | | | | |
|---|---|---|---|---|---|---|---|---|
| 4f (HbT max peak, 10-30s) | WT: 9.602, AD: 8.250, ATH: 8.731, MIX: 9.128 | WT: 5.815, AD: 5.357, ATH: 5.257, MIX: 6.230 | Linear mixed model (fixed factor: disease group) | F=0.1708 | 4.2858 | 3 | 0.915 | AD-ATH p=0.9999, AD-MIX p=0.9927, AD-WT p=0.9310, ATH-MIX p=0.9941, ATH-WT p=0.9213, MIX-WT p=0.9919 |
| 4g (HbR AUC, 10-30s) | WT: -11.18, AD: -8.719, ATH: -7.927, MIX: -8.306 | WT: 18.61, AD: 11.58, ATH: 14.80, MIX: 16.04 | Linear mixed model (fixed factor: disease group) | F=0.3598 | 74.169 | 3 | 0.7825 | AD-ATH p=0.9936, AD-MIX p=0.9988, AD-WT p=0.9247, ATH-MIX p=0.9998, ATH-WT p=0.7551, MIX-WT p=0.8796 |
| 2h (Locomotion AUC, 10-30s) | WT: 24.02, AD: 9.038, ATH: 17.88, MIX: 32.93 | WT: 38.65, AD: 10.79, ATH: 28.51, MIX: 59.68 | Linear mixed model (fixed factor: disease group) | F=0.9565 | 840.14 | 3 | 0.4295 | AD-ATH p=0.8595, AD-MIX p=0.3891, AD-WT p=0.6554, ATH-MIX p=0.7209, ATH-WT p=0.9606, MIX-WT p=0.9376 |

**SR5: Figure 5: No differences in performance on a novel object recognition task between disease groups.** For the distance run during training and testing (5b), velocity during training and testing (5c), and the preference index (5d) (dependent variables) the data was compared across disease groups (independent variable, 4 levels: WT, AD, ATH, MIX). All datasets were first tested for normality using the Shapiro-Wilks test (e.g. Figure 5b distance run the following individual groups were each tested: training WT, testing WT, training AD, testing AD, training ATH, testing ATH, training MIX, testing MIX), and for variance between groups using the Brown-Forsythe test (distance run training p=0.98, distance run testing p=0.92, velocity training p=0.75, velocity testing p=0.88, preference index p=0.95), with non-significant values indicating the assumptions of the one-way ANOVAs had been met. We saw no significant differences in performance on the novel object recognition task between disease groups. Due to the AD mice performing below chance for the preference index (5d) we have also displayed the Tukey post-hoc comparisons to highlight we still detect no differences between the AD mice or any other mouse groups.

| **Figure** | **Mean** | **SD** | **Test** | **Test Statistic** | **Degrees of freedom** | **P Value** |
|---|---|---|---|---|---|---|

| | | | | | | |
|---|---|---|---|---|---|---|
| 5b (distance run, cm) | Training:<br>WT: 2437, AD: 2482, ATH: 2815, MIX: 2828<br>Testing:<br>WT: 1982, AD: 2057, ATH: 2239, MIX: 2083 | Training: WT: 581.5, AD: 390.6, ATH: 531.6, MIX: 546.6<br>Testing:<br>WT: 492.2, AD: 484, ATH: 752.9, MIX: 761.7 | One-way ANOVA | Training: F=1.107<br>Testing:<br>F=0.2214 | DFn=3, DFd=24 | Training: p=0.3657<br>Testing: p=0.8805 |
| 5c (velocity, cm/s) | Training:<br>WT: 4.358, AD: 4.612, ATH: 5.098, MIX: 4.85<br>Testing:<br>WT: 3.539, AD: 3.571, ATH: 4.00, MIX: 3.752 | Training: WT: 1.083, AD: 1.353, ATH: 1.137, MIX: 0.9114<br>Testing:<br>WT: 1.067, AD: 0.996, ATH: 1.333, MIX: 1.487 | One-way ANOVA | Training: F=0.5918<br>Testing:<br>F=0.2293 | DFn=3, DFd=24 | Training: p=0.6264<br>Testing: p=0.8751 |
| 5d (preference index) | WT: 58.2, AD: 48.94, ATH: 61.46, MIX: 56.95 | WT: 17.06, AD: 13.49, ATH: 13.12, MIX: 17.42 | One-way ANOVA | F=0.8800 | DFn=3, DFd=24 | p=0.4653 |
| | Tukey posthoc comparison: WT-AD p=0.7129; WT -ATH p=0.9744; WT-MIX p=0.9989; AD-ATH p=0.3909; AD-MIX p=0.7934; ATH-MIX p=0.9367 | | | | | |

**SR6: Figure 6: No differences in pathology across disease models.** To assess pathology between our disease models we compared amyloid plaque coverage (% area) in AD and MIX mice, and aortic arch plaque load in ATH and MIX mice. The Shapiro-Wilks test of normality and Brown-Forsythe or F test of independence were conducted for each group to check the t-test or ANOVA assumptions were met, and all contained data which was normally distributed (6b: ATH: p=0.74, MIX: p=0.45; 6d: AD cortex p=0.73, MIX cortex p=0.62, AD hippocampus p=0.87, MIX hippocampus p=0.10) and samples independent (6b p=0.25; 6d p=0.40). As the aortic arch comparison had only two levels (independent variable disease group, 2 levels: ATH, MIX) an unpaired t-test was conducted; whereas for the amyloid beta assessment a mixed experimental design was used (disease 2 levels: AD, MIX; brain region 2 levels: cortex, HC) meaning a mixed ANOVA was conducted.

| Figure | Mean | SD | Test | Test Statistic | Degrees of freedom | P Value |
|---|---|---|---|---|---|---|
| 6b (aortic plaque load) | ATH: 18.26<br><br>MIX: 17.75 | ATH: 4.788<br><br>MIX: 2.287 | Unpaired t-test | t=0.1977 | DFn=7, DFd=3 | p=0.2519 |
| 6d (Abeta plaque load) | AD Cortex: 2.118<br><br>MIX Cortex: 1.489<br><br>AD HC: 2.271<br><br>MIX HC: 1.332 | AD Cortex: 0.8311<br><br>MIX Cortex: 0.7746<br><br>AD HC: 0.3132<br><br>MIX HC: 0.5416 | Mixed ANOVA | Genotype F=3.3470, Brain region F=0.00013, genotype * brain region F=0.7450 | DFn 1, DFd 7 | Genotype p=0.110, Brain region p=0.991, genotype * brain region p=0.417 |
